# Metabolic Perceptrons for Neural Computing in Biological Systems

**DOI:** 10.1101/616599

**Authors:** Amir Pandi, Mathilde Koch, Peter L Voyvodic, Paul Soudier, Jerome Bonnet, Manish Kushwaha, Jean-Loup Faulon

## Abstract

Synthetic biological circuits are promising tools for developing sophisticated systems for medical, industrial, and environmental applications. So far, circuit implementations commonly rely on gene expression regulation for information processing using digital logic. Here, we present a new approach for biological computation through metabolic circuits designed by computer-aided tools, implemented in both whole-cell and cell-free systems. We first combine metabolic transducers to build an analog adder, a device that sums up the concentrations of multiple input metabolites. Next, we build a weighted adder where the contributions of the different metabolites to the sum can be adjusted. Using a computational model trained on experimental data, we finally implement two four-input “perceptrons” for desired binary classification of metabolite combinations by applying model-predicted weights to the metabolic perceptron. The perceptron-mediated neural computing introduced here lays the groundwork for more advanced metabolic circuits for rapid and scalable multiplex sensing.

## Introduction

Living organisms are information-processing systems that integrate multiple input signals, perform computations on them, and trigger relevant outputs. The multidisciplinary field of synthetic biology has combined their information-processing capabilities with modular and standardized engineering approaches to design sophisticated sense-and-respond behaviors^1–3^. Due to similarities in information flow in living systems and electronic devices^4^, circuit design for these behaviors has often been inspired by electronic circuitry, with substantial efforts invested in implementing logic circuits in living cells^4–6^. Furthermore, synthetic biological circuits have been used for a range of applications including biosensors for detection of pollutants^7, 8^ and medically-relevant biomarkers^9, 10^, smart therapeutics^11, 12^, and dynamic regulation and screening in metabolic engineering^13, 14^.

Synthetic circuits can be implemented at different layers of biological information processing, such as: (i) the genetic layer comprising transcription^15^ and translation^16^, (ii) the metabolic layer comprising enzymes^17, 18^, and (iii) the signal transduction layer comprising small molecules and their receptors^19, 20^. Most designs implemented thus far have focused on the genetic layer, developing circuits that perform computations using elements such as feedback control^21^, memory systems^22, 23^, amplifiers^24, 25^, toehold switches^26^, or CRISPR machinery^27, 28^. However, gene expression regulation is not the only way through which cells naturally perform computation. In nature, cells carry out parts of their computation through metabolism, receiving multiple signals and distributing information fluxes to metabolic, signaling, and regulatory pathways^17, 29, 30^. Integrating metabolism into synthetic circuit design can expand the range of input signals and communication wires used in biological circuits, while bypassing some limitations of temporal coordination of gene expression cascades^31, 32^.

The number of inputs processed by synthetic biological circuits has steadily increased over the years, including physical inputs like heat, light, and small molecules such as oxygen, IPTG, aTc, arabinose and others^21, 33–36^. However, most of these circuits process input signals using digital logic, which despite its ease of implementation lacks the power that analog logic can offer^1, 37, 38^. The power of combining digital and analog processing is exemplified by the “perceptron”, the basic block of artificial neural networks inspired by human neurons^39^ that can, for instance, be trained on labelled input datasets to perform binary classification. After the training, the perceptron computes the weighted sum of input signals (analog computation) and makes the classification decision (digital computation) after processing it through an activation function.

Here we describe the development of complex metabolic circuitry implemented using analog logic in whole-cell and cell-free systems by means of enzymatic reactions. For circuit design, we first employ computational design tools, Retropath^40^ and Sensipath^41^, that use biochemical retrosynthesis to predict metabolic pathways and biosensors. We then build and model three whole-cell metabolic transducers and an analog adder to combine their outputs. Next, we transfer our metabolic circuits to a cell-free system^42, 43^ in order to take advantage of the higher tunability and the rapid characterization it offers^44–46^, expanding our system to include multiple weighted transducers and adders. Finally, using our integrated model trained on the cell-free metabolic circuits we build a more sophisticated device called the “metabolic perceptron”, which allows desired binary classification of multi-input metabolite combinations by applying model-predicted weights on the input metabolites before analog addition, and demonstrate its utility through two examples of four-input binary classifiers. Altogether, in this work we demonstrate the potential of synthetic metabolic circuits, along with model-assisted design, to perform complex computations in biological systems.

## Results

### Whole-cell processing of hippurate, cocaine and benzaldehyde inputs

To identify the metabolic circuits to build, we use our metabolic pathway design tools, Retropath^40^ and Sensipath^41^. These tools function using a set of sink compounds at the end of a metabolic pathway, here metabolites from a dataset of detectable compounds^47^, and a set of source compounds that can be used as desired inputs for the circuit. The tools then propose pathways and the enzymes that can catalyze the necessary reactions, allowing for promiscuity. Our metabolic circuit layers are organized according to the main processing functions: transduction and actuation (**Figure 1a**). Transducers are the simplest metabolic circuits that function as sensing enabling metabolic pathways (SEMP)^48^, consisting of one or more enzymes that transform an input metabolite into a transduced metabolite. The transduced molecule, in turn, is detected through an actuation function that is implemented using a transcriptional regulator.

**Figure 1.**
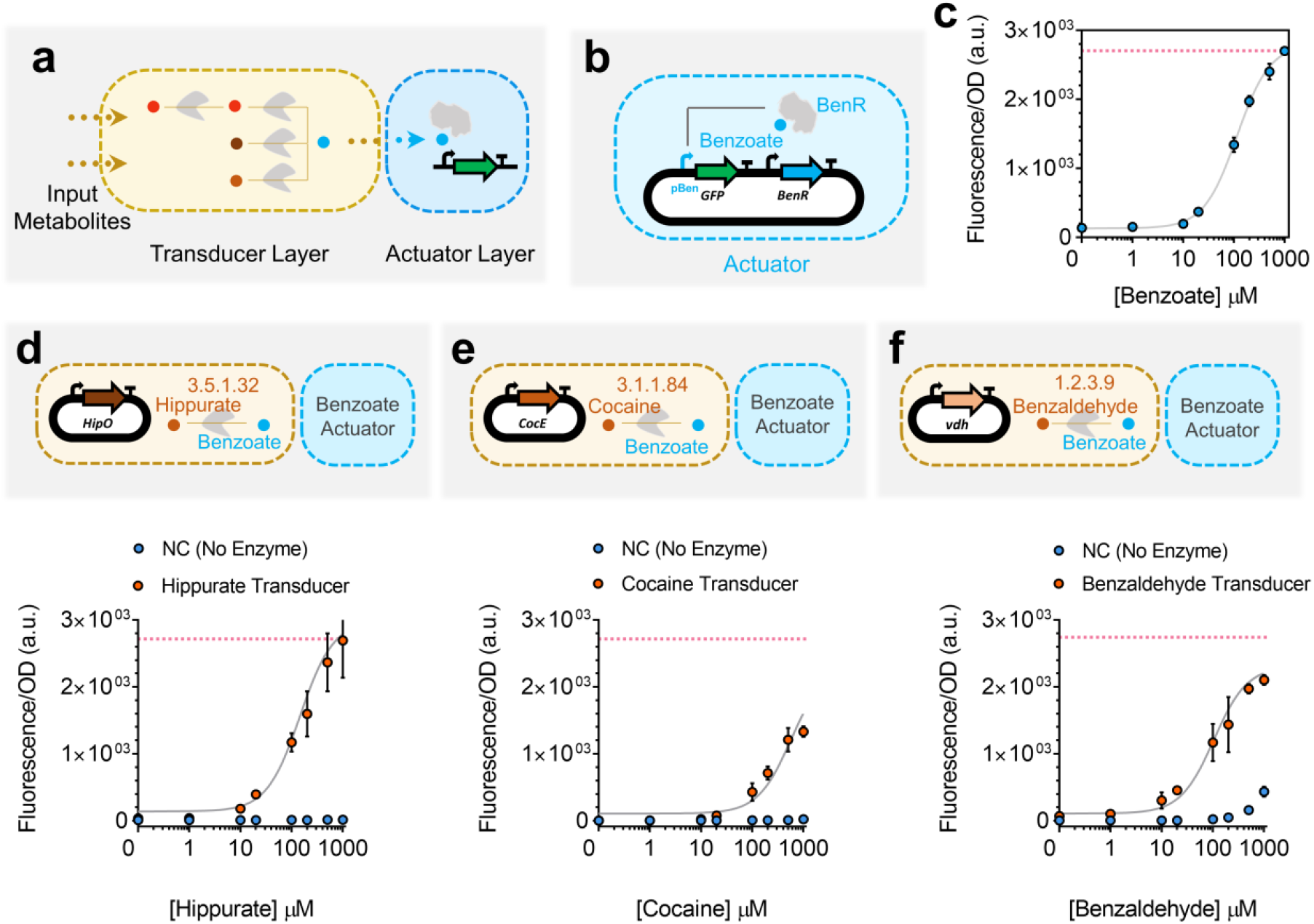
Whole-cell actuator and metabolic transducers. (**a**) Designed synthetic metabolic circuits using Retropath40 or Sensipath41 consist of a transducer layer and an actuator layer. (**b**) Open-loop circuit construction of the benzoate actuator, which is used downstream of transducer metabolic circuits in this work. For the open-loop circuit, the transcription factor (TF) is expressed constitutively under control of the promoter J23101 and RBS B0032. (**c**) Dose-response plot of the open-loop circuit for the benzoate actuator. The gray curve is a model-fitted curve (see Methods section) for the open-loop circuit. (**d,e,f**) Whole-cell metabolic transducers for hippurate (**d**), cocaine (**e**) and benzaldehyde (**f**) represented in dose-response plots (orange circles) and their associated dose-response when there is no enzyme present (blue circles). The red dotted lines refer to the maximum signal from the actuator (**c**). The transducer output benzoate is reported through the open-loop circuit actuator. The enzymes are expressed under constitutive promoter J23101 and RBS B0032. All data points and the error bars are the mean and standard deviation of normalized values from three measurements.

We used benzoate as our transduced metabolite, its associated transcriptional activator BenR, and the responsive promoter pBen to construct the actuator layer of our whole-cell metabolic circuits^49^. To compare the shape of the response curve, we constructed the actuator layer in two formats: (i) an open-loop circuit (**Figure 1b**) and (ii) a feedback-loop circuit (**Figure S1**). When compared to the open-loop format, the feedback-loop circuit has previously been shown to exhibit linear dose-response to input^21, 50^. We found that while the feedback-loop format does linearize the actuator response curve, apparent toxicity at high benzoate concentrations reduces the usable activator dynamic range (**Figure S1**). Therefore, we selected the open-loop format due to its higher dynamic range of activation (**Figure 1c**), setting the maximum concentration of benzoate used in this work to the saturation point of this open-loop circuit.

Building on our previous work^48^, we next implemented three upstream transducers that convert different input metabolites into benzoate for detection by the actuator layer already tested. The transducer layers were composed of enzymes HipO for hippurate (**Figure 1d**), CocE for cocaine (**Figure 1e**), and vdh for benzaldehyde (**Figure 1f**). Compared to the benzoate output signal, we found that the transduction capacities of the three transducers were 99.6%, 49.2%, and 77.8%, respectively (**Supplementary Figure S2**), indicating a partial dissipation in signal.

### A Whole-cell metabolic concentration adder

A metabolic concentration adder is a device composed of more than one transducer that converts their respective input metabolites into a common transduced output metabolite. For our whole-cell concentration adder, we combined two transducers to build a hippurate-benzaldehyde adder actuated by the benzoate circuit (**Figure 2a**). Unlike digital bit-adders that exhibit an ON-OFF digital behavior, our metabolic adders exhibit a continuous analog behavior that is natural for metabolic signal conversion^51^ (**Figure 2b** and **Supplementary Figure S3**). Increasing the concentration of one of the inputs at any fixed concentration of the other shows an increase in the output benzoate, and thus in the resulting fluorescence (**Figure 2b** and **Supplementary Figure S3**).

**Figure 2.**
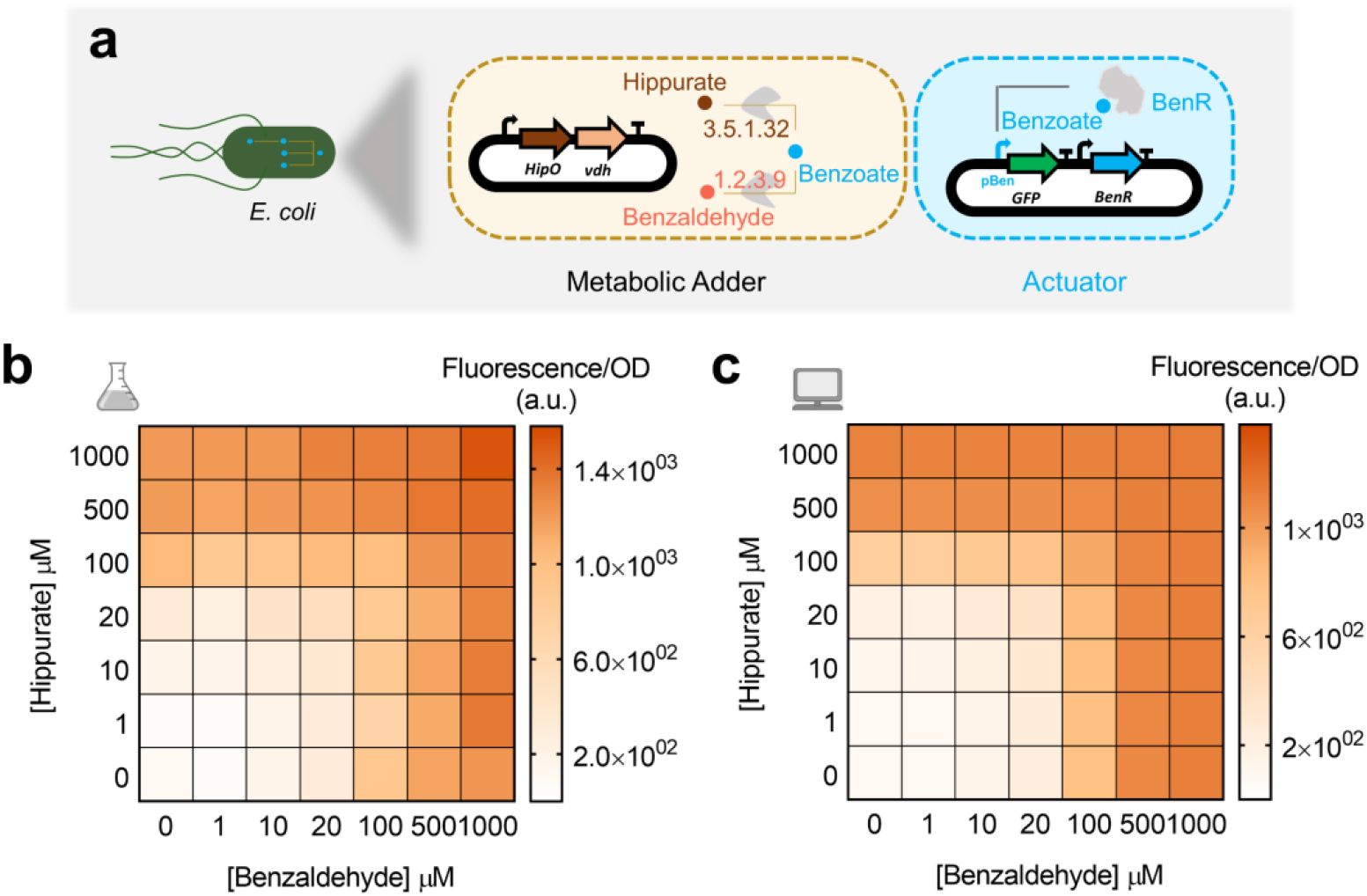
Whole-cell metabolic adder of hippurate and benzaldehyde. (**a**) Hippurate and benzaldehyde transducers are combined to build a metabolic adder producing a common output, benzoate, which is reported through the benzoate actuator. The enzymes are expressed in one operon under control of constitutive promoter J23101 and RBSs B0032 for HipO and B0034 for vdh. (**b**) Heatmap representing the output of the adder while increasing the concentration of both inputs, hippurate and benzaldehyde. All data points are the mean of normalized values from three measurements. (**c**) Model simulations for experimental conditions presented in (**b**). The model was fitted on transducer data and resource competition data.

The maximum output signal for our adder, when hippurate and benzaldehyde were both at the maximum concentration of 1000 µM, was lower than the maximum signal produced by hippurate and benzaldehyde transducers alone (**Supplementary Figure S2**). However, as seen above, the difference between the maximum signal of their transducers and the actuator was smaller. This dissipation in signal from the transducers to the adders and from the actuators to the transducers (**Supplementary Figure S2**) could either be because of resource competition (as a result of adding more genes) or because of enzyme efficiency (as a result of poorly balanced enzyme stoichiometries). To test these two hypotheses, we investigated the effect of the enzymes on cellular resource allocation. For this purpose, the cocaine transducer and the hippurate-benzaldehyde adder were characterized by adding benzoate to these circuits (**Supplementary Figures S4** and **S5**). Comparing the results of these characterizations with the benzoate actuator reveals that dissipation in signal from the transducers to the adders is due to resource competition, whereas that from the actuators to the transducers is due to enzyme efficiency.

In order to gain quantitative understanding of the circuits’ behavior, we empirically modeled their individual components to see if we were able successfully capture their behavior. We first modeled the actuator (gray curve in **Figure 1c**) using Hill formalism^52^ as it is the component that is common to all of our outputs and therefore constrains the rest of our system. We then modeled our transducers, considering enzymes to be modules that convert their respective input metabolites into benzoate, which is then converted to the fluorescence output already modeled above. This simple empirical modeling strategy reproduces our transducer data (results not shown). To incorporate observations made in **Supplementary Figure S4** and **S5**, we included resource competition in our models to explain circuits with one or more transducers. To this end, we extended the Hill model to account for resource competition following previous works^53, 54^, with a fixed pool of available resources for enzyme and reporter protein production that is depleted by the transducers. This extension is further presented in the Methods section. We trained our model on all transducers, with and without resource competition (i.e. individual transducers, or transducers where another enzyme competes for the resources). This model (presented in gray lines in **Figure 1d,e,f** and **Figure 2c**), which was not trained on adder data but only on actuator, transducer, and transducers with resource competition data, recapitulates it well. This indicates that the model accounts for all important effects underlying the data. The full training process is presented in the Methods section, and a table summarising scores of estimated goodness of fit of our model is presented in **Supplementary Table S1**.

### Cell-free processing of multiple metabolic inputs

Cell-free systems have recently emerged as a promising platform^42^ that provide rapid prototyping of large libraries by serving as an abiotic chassis with low susceptibility to toxicity. We took advantage of an *E. coli* cell-free system with the aim of increasing the computational potential of metabolic circuits in several ways (**Figure 3a**). Firstly, a higher number of genes can be simultaneously and combinatorially used to increase the complexity and the number of inputs for our circuits. Secondly, the lower noise provided by the absence of cell growth and maintenance of cellular pathways^55^ improves the predictability and accuracy of the computation. Thirdly, having genes cloned in separate plasmids enables independent tunability of circuit behavior by varying the concentration of each part individually. Finally, cell-free systems are highly adjustable for different performance parameters and components. In all, these advantages of cell-free systems enable us to develop more complex computations than the whole-cell adder.

**Figure 3.**
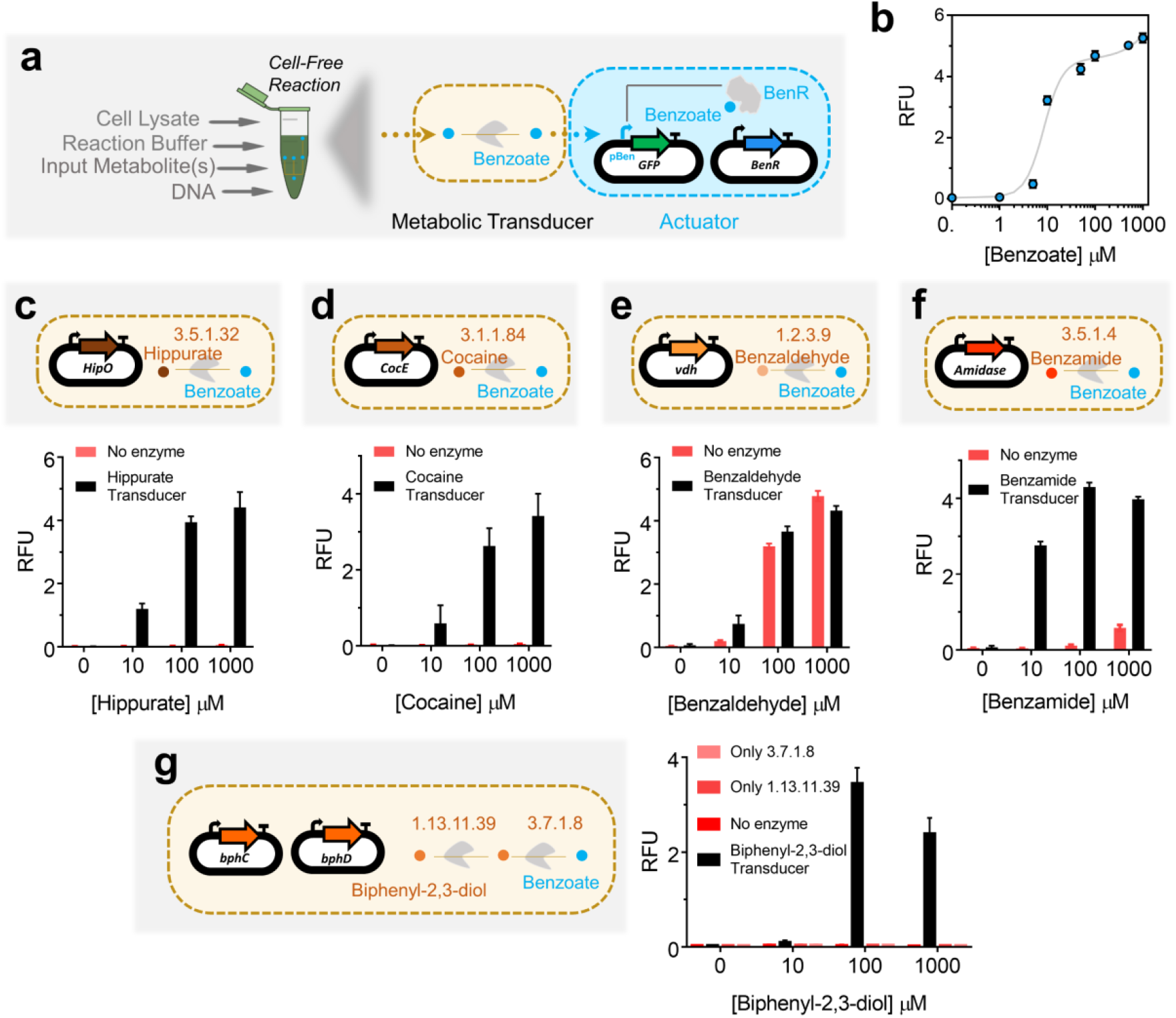
Cell-free actuator and metabolic transducers. (**a**) Implementing benzoate actuator and transducers in *E. coli* transcription/translation (TXTL) cell-free system. Cell-free reactions are composed of cell lysate, reaction buffer (energy source, tRNAs, amino acids, etc.) and DNA plasmids. (**b**) Dose-response plot of the benzoate actuator in the cell-free system with 30 nM of TF-plasmid (constitutively expressed BenR) and 100 nM of reporter plasmid (pBen-sfGFP) per reaction. The data points represent the dose-response of the actuator to different concentrations of benzoate and the gray curve is a model-fitted curve on actuator data (**c,d,e,f,g**). Cell-free transducers coupled with the benzoate actuator for hippurate (**c**), cocaine (**d**), benzaldehyde (**e**), benzamide (**f**), and biphenyl-2,3-diol (**g**), which is composed of two enzymes. All enzymes are cloned in a separate plasmid under the control of a constitutive promoter J23101 and RBS B0032. 10 nM of each plasmid was added per reaction. The bars are the response of the circuits to different concentrations of input with (transducers, black bars) and without enzyme (red bars). All data are the mean and the error bars are the standard deviation of normalized values from three measurements (RFU: Relative Fluorescence Unit).

Following from our recent work^56^, we first characterized a cell-free benzoate actuator to be used downstream of other metabolic transducers. **Figure 3a** shows a schematic of the cell-free benzoate actuator composed of a plasmid encoding the BenR transcriptional activator and a second plasmid expressing sfGFP reporter under the control of a pBen promoter. This actuator showed a higher operational range than the whole-cell counterpart (**Figure 1c**). The optimal concentration of the TF plasmid (30 nM) and the reporter plasmid (100 nM) were taken from our recent study^56^. Following successful implementation of the actuator, we proceeded to build five upstream cell-free transducers for hippurate, cocaine, benzaldehyde, benzamide, and biphenyl-2,3-diol (**Figure 3c,d,e,f,g**) that convert these compounds to benzoate. Each of the five transducers used 10 nM of enzyme DNA per reaction, except the biphenyl-2,3-diol transducer that used two metabolic enzymes with 10 nM DNA each.

Compared to its whole-cell counterpart (**Figure 1f**), in the cell-free transducer reaction (**Figure 3e**) benzaldehyde appears to spontaneously oxidise to benzoate without the need of the transducer enzyme vdh. This behavioral difference between the whole-cell and cell-free setups could be due to the difference in redox states inside an intact cell and the cell-free reaction mix^57, 58^. Furthermore, benzamide and biphenyl-2,3-diol transducers exhibit inhibition in fluorescence outputs at very high (1000 μM) input concentrations.

### Cell-free weighted transducers and adders

After characterizing different transducers in the cell-free system that enable building a multiple-input metabolic circuit, we sought to rationally tune the transducers. Cell-free systems allow independent tuning of each plasmid by pipetting different amounts of DNA. We applied this advantage to weight the flux of enzymatic reactions in cell-free transducers (**Figure 4a**). The concentration range we used was taken from our recent study^56^, in order to have an optimal expression with minimum resource competition. We built four weighted transducers for hippurate (**Figure 4b**), cocaine (**Figure 4c**), benzamide (**Figure 4d**) and biphenyl-2,3-diol (**Figure 4e**). Increasing the concentration of the enzymes produces a higher amount of benzoate from the input metabolites, and hence higher GFP fluorescence. Compared to the others, the hippurate transducer reached higher GFP expression at a given concentration of the enzyme and the input, and biphenyl-2,3-diol reached the weakest signal. For the biphenyl-2,3-diol transducer built with two enzymes (**Figure 4e**), both enzymes are added at the same concentration (e.g., 1 nM of “enzyme DNA” indicates 1 nM each of plasmids encoding enzymes bphC and bphD).

**Figure 4.**
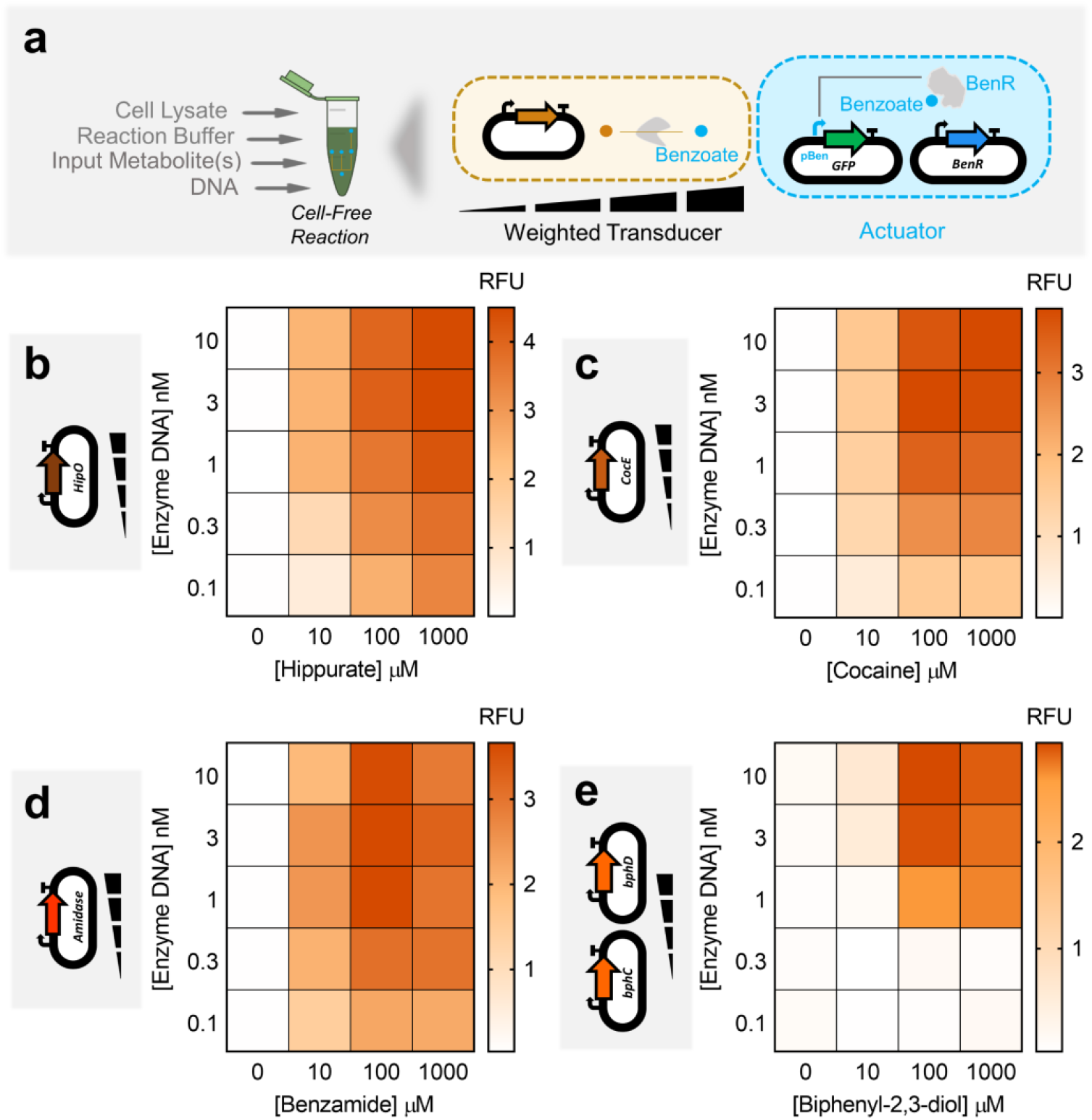
Cell-free weighted transducers characterized by varying the concentration of the enzyme DNA. (**a**) In the cell-free system, the circuits can be tuned by varying the amount of each enzyme pipetted per reaction. Weighted transducers are characterized by varying the concentration of the enzymes in transducers which then are reported through the benzoate actuator. The range of the concentrations was varied to get optimal expression and minimum resource competition. (**b,c,d,e**) Heatmaps representing weighted transducers at different concentrations of input molecules and enzymes DNA for hippurate (**b**), cocaine (**c**), benzamide (**d**) and biphenyl-2,3-diol (**e**). For the biphenyl-2,3-diol weighted transducer (**e**), concentrations represent those of each metabolic plasmid (e.g., 1 nM of “enzyme DNA” refers to 1 nM of bphC plus 1 nM of bphD). See **Supplementary Figure S6** for model results of each weighted transducer. All data are the mean of normalized values from three measurements. (RFU: Relative Fluorescence Unit).

Data in **Figure 4** show that similar output levels can be achieved for different input concentrations, provided the appropriate transducer concentrations are used. In the next step, we applied this finding to build hippurate-cocaine weighted adders by altering either the concentration of the enzymes or the concentration of the inputs (**Figure 5a**). The fixed-input adder is an adder in which the concentration of inputs, hippurate and cocaine, are fixed to 100 µM and the concentration of the enzymes is altered (top panel in **Figure 5b**). In this device, the weight of the reaction fluxes is continuously tunable. We then characterized a fixed-enzyme adder by fixing the concentration of the enzymes (1 nM for HipO, 3 nM for CocE; the cocaine signal is weaker, which is why a higher concentration of its enzyme is used) and varying the inputs, hippurate and cocaine (top panel in **Figure 5c**).

**Figure 5.**
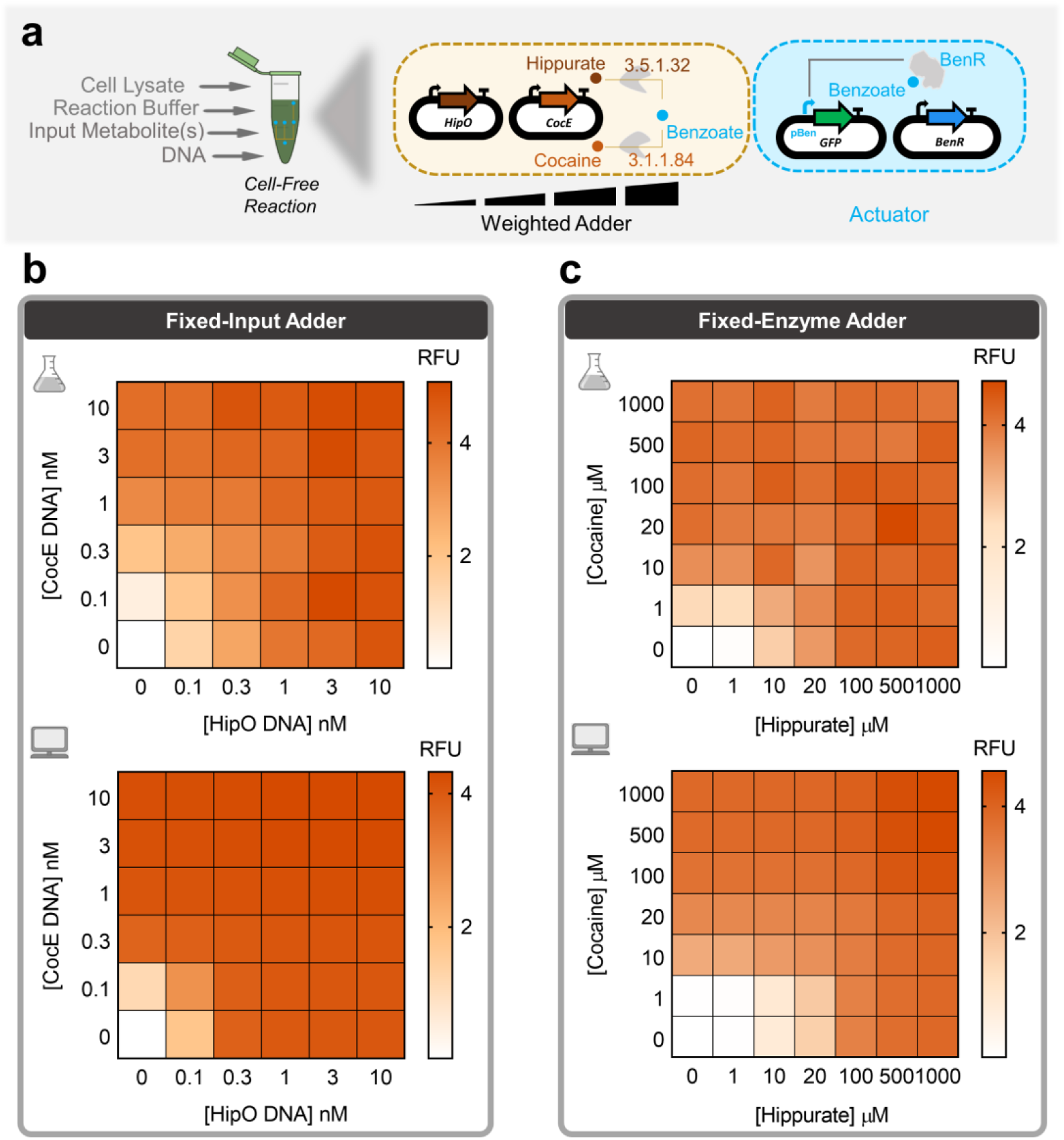
Multiple transducers are combined to shape an adder while weighing inputs or enzymes. (**a**) Cell-free adder characterization by varying the concentration of either inputs or enzymes producing different levels of fluorescence through the actuator. (**b**) Heatmap showing fixed-input adder in which the inputs, hippurate and cocaine, are fixed to 100 µM and concentrations of associated enzyme are altered by altering the concentration of plasmid DNA encoding them. *Top:* Cell-free experiment of hippurate-cocaine fixed-input (weighted) adder. *Bottom:* Model simulation (prediction) of hippurate-cocaine fixed-input (weighted) adder. (**c**) Fixed-enzyme adder with fixed concentrations of the enzyme DNAs, 1 nM for HipO and 3 nM for CocE, and various concentrations of the inputs, hippurate and cocaine. *Top:* Cell-free experiment of hippurate-cocaine fixed-enzyme adder. *Bottom:* Model simulations (prediction) of hippurate-cocaine fixed-enzyme adder. All data are the mean of normalized values from three measurements. (RFU: Relative Fluorescence Unit).

In order to have the ability to build any weighted adder with predictable results, we developed a model that accounts for the previous data. We first empirically modeled the actuator (gray curve in **Figure 3b**) since all other functions are constrained by how the actuator converts metabolite data (benzoate) into a detectable signal (GFP). We then trained our model with individual weighted transducers (**Supplementary Figure S6**) and predicted the behaviors of the weighted adders (bottom panel in **Figure 5b,c**). The results shown in **Figure 5b,c** indicate that our model describes the adders well, despite being trained only on transducer data. **Supplementary Table S2** summarizes the different scores to estimate goodness of fit of our model. Briefly, the model quantitatively captures the data but tends to overestimate values at intermediate enzyme concentration ranges and does not capture the inhibitory effect observed at the high concentration of benzamide or biphenyl-2,3-diol, as this was not accounted for in the model.

Using the above strategy, we can build any weighted adder for which we have pre-calculated the weights using the model on weighted transducers. We use this ability in the following section to perform more sophisticated computation for a number of classification problems.

### Cell-free perceptron for binary classifications

The perceptron algorithm was first developed to computationally mimic the neuron’s ability to process information, learn, and make decisions^59^. Perceptrons are the basic blocks of artificial neural networks enabling the learning of deep patterns in datasets by training the model’s input weights^60^. Like a neuron, the perceptron receives multiple input signals (*X_i_*) and triggers an output depending on the weighted (*W_i_*) sum of the inputs^39^. A perceptron can be used to classify a set of input combinations after it is trained on labeled data. In binary classification, the weighted sum is first calculated (Σ*W_i_.X_i_*) and an activation function (*f*), coupled with a decision threshold d, finally makes the decision: ON if f(Σ*W_i_.X_i_)* > d, OFF otherwise (**Figure 6a**). The activation function could be linear or non-linear (Sigmoid, tanh, ReLU, etc.) depending on the problem^61^, although a sigmoid is generally used for classification.

**Figure 6.**
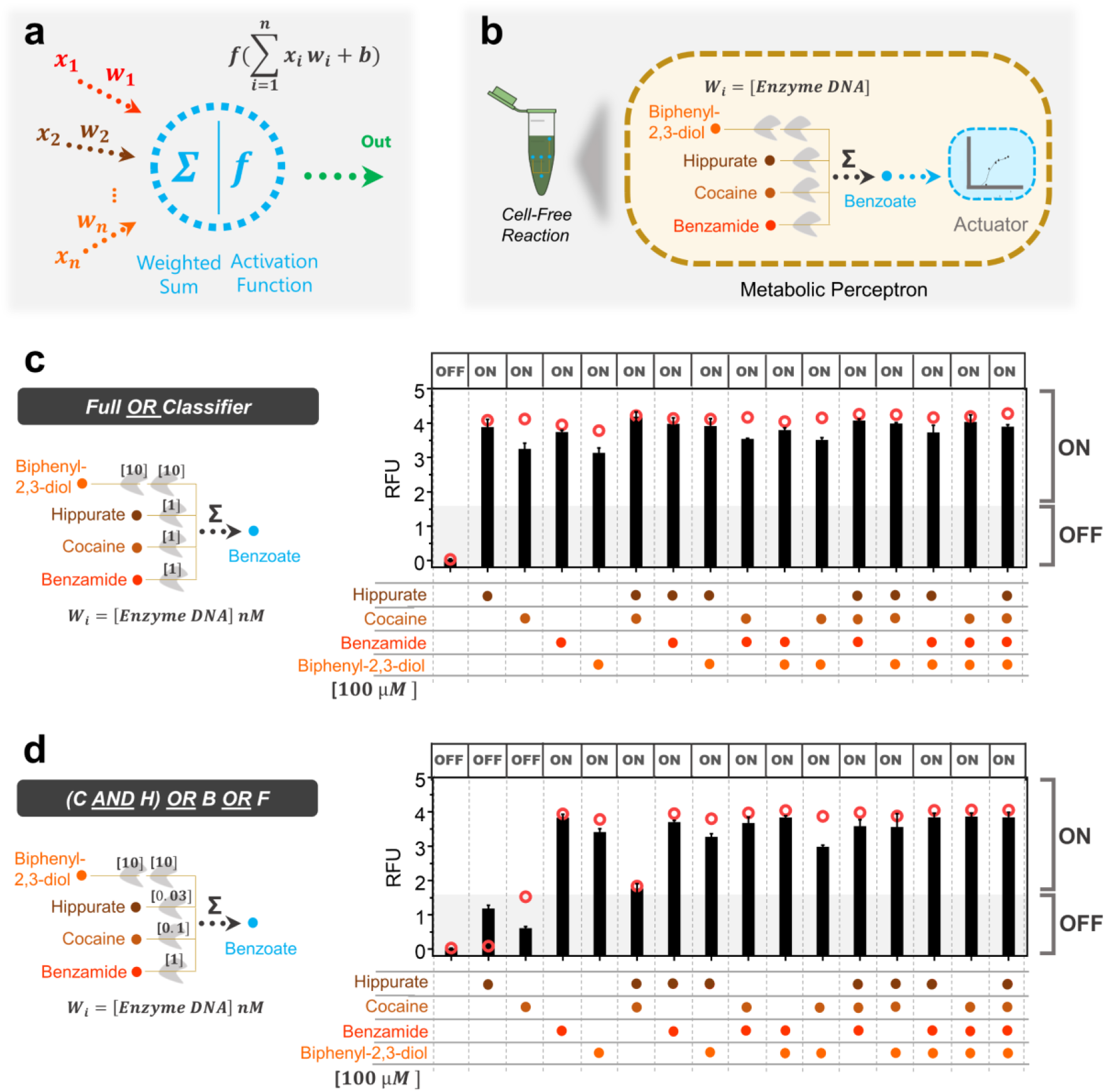
Cell-free perceptron enabling development of classifiers. (**a**) A perceptron scheme showing the inputs and their associated weights, the computation core, and the output. The perceptron computes the weights and actuates the weighted sum through an activation function. (**b**) Metabolic perceptron integrating multiple inputs and actuating an output. The benzoate actuator acts as the activation function of the perceptron reporting the sum of benzoate produced by the metabolic perceptron. Hippurate, cocaine, benzamide, and biphenyl-2,3-diol are the inputs of the metabolic perceptron fixed to 100 µM. The weights of the perceptron are the concentration of the enzymes calculated using the model made on weighted metabolic circuits (red circles). These weights are calculated to develop two classifiers using the metabolic perceptron and benzoate actuator. “Full OR” classifier (**c**), “[cocaine *(C)* AND hippurate *(H)*] OR benzamide *(B)* OR biphenyl-2,3-diol *(F)*” classifier (**d**) are the two classifiers built using this metabolic perceptron. The “Full OR” classifier (**c**) classifies to “OFF” when none of the inputs is present and it passes an arbitrary threshold to “ON” when any of the inputs or their combinations are present. The second classifier (**d**) performs a more complex computation. The shading represents the arbitrary threshold that allows for perceptron decision making and the panel of “OFF” and “ON” at the top of the bars are the expected output of the classifiers. All data are the mean and the error bars are the standard deviation of normalized values from three measurements and red circles are the model predictions. (RFU: Relative Fluorescence Unit).

Since our weighted transducer models have already been trained on the cell-free experimental data, we checked if we could use them to calculate the weights needed to classify different combinations of two inputs: hippurate and cocaine. We tested our model on five different binary classification problems, A to E (**Supplementary Figure 7**). For each problem, the two types of data were represented as a cluster of dots on the scatter plot. The trained model was then used to identify weights needed to be applied to the weighted transducers such that a decision threshold ‘d’ exists to classify the two clusters into red (ON, >d) or blue (OFF, <= d). The lines shown in **Supplementary Figure 7** plots show three iso-fluorescence lines that represent the threshold that classifies the data into the binary categories: ON and OFF. These theoretical classification problems demonstrate the ability of our trained perceptron model to successfully carry out binary classification.

Using the integrated model from our weighted transducers and adders, we next sought to design four-input classifiers using a metabolic perceptron, and test them experimentally. Our metabolic perceptron is a device enabling signal integration of multiple inputs with associated weights, represented by enzyme DNA concentrations (**Figure 6b**). The 4-input adder performs the weighted sum and the benzoate actuator acts as the activation function of the metabolic perceptron. The weights can be adjusted to implement different classification functions. To illustrate the potential of building perceptrons with metabolic weighted adders, we computed adder weights using our model for two different classifiers: a simple classifier equivalent to a “full OR” gate (**Figure 6c**), and a more complex classifier equivalent to a “[cocaine AND hippurate] OR benzamide OR biphenyl-2,3-diol” gate (**Figure 6d**). Weight calculation methods are reported in the Methods section.

For the classifiers, the input metabolites are fixed to 100 µM, as it allows the best ON-OFF behavior for all inputs and weight-tuning according to model simulations (results not shown). The model accurately predicted weights to obtain the simple “full OR” classifier behavior (**Figure 6d**), as well as cocaine, benzamide, and biphenyl-2,3-diol weights for the second complex classifier. The initial weights computed by the model are presented in **Supplementary Figure S8**. The optimal weight of HipO (hippurate transducing enzyme) was calculated to be 0.1 nM, which leads to higher signals than predicted, particularly for the “ON” behavior with only hippurate. To further characterize the HipO weights at still lower concentrations of the enzyme, we performed an additional complementary characterization (**Supplementary Figure S9**). Our aim here was to find a weight for HipO through which a classifier outputs a low signal (“OFF”) with only hippurate and high signal (“ON”) when coupled with other inputs. We arrived at 0.03 nM HipO which exhibited this shifting behavior between “OFF” and “ON” (**Figure 6d** and **Supplementary Figure S9**). Using our model-guided design and rapid cell-free prototyping on the HipO weight, we were able to design two 4-input binary classifiers. In **Figure 6c,d** red circles are the weights predicted with 0.03 nM for HipO and the bars are experimental results. All actual values of the model and the experiments are provided in **Supplementary Table S7**.

## Discussion

Computing in synthetic biological circuits has largely relied on digital logic-gate circuitry for almost two decades^5, 62^, treating inputs as either absent (0) or present (1). While such digital abstraction of input signals provides conceptual modularity for circuit design, it is less compatible with the physical-world input signals that vary between low and high values on a continuum^37^. As a result, digital biological circuits must carefully match input-output dynamic ranges at each layer of signal transmission to ensure successful signal processing^2, 30^. More recently, the higher efficiency of analog computation on continuous input has been recognized^63^, and some analog biological circuits have started emerging^21^. In this regard, using metabolic pathways for cellular computing seems like a natural progression for analog computation in biological systems^21, 30^.

In this study, we investigated the potential of metabolism to perform analog computations using synthetic metabolic circuits. To that end, we first established a benzoate actuator to report the output from our metabolic circuits in both whole-cell and cell-free systems (**Figures 1c and 3b**). Upstream of the actuator, we constructed hippurate, cocaine, and benzaldehyde transducers in the whole-cell system (**Figures 1d,e,f**) and a metabolic adder by combining the benzaldehyde and hippurate transducers (**Figure 2**). Similarly, we constructed hippurate, cocaine, benzaldehyde, benzamide, and biphenyl-2,3-diol transducers in the cell-free system (**Figures 3c,d,e,f,g**) and weighted adders by combining them (**Figure 5**). Compared to the numerous digital biological devices, which compute through multi-layered genetic logic circuits, the metabolic adder is a simple one-layered device with fast execution times.

Our computational models trained only on the actuator and transducer data predicted adder behaviors with high accuracy (**Supplementary Tables S1 and S2**). This further enabled us to calculate the required weights for more complex “metabolic perceptrons” that compute weighted sums from multiple inputs and use them to classify the multi-input combinations in a binary manner (**Figures 6 and S7**). Although we used fixed concentrations of inputs to demonstrate the ability of our perceptrons to classify, models trained on characterization data from weighted transducers should enable one to build classifiers for other concentrations in the operational range of the transducers (**Supplementary Figure S10**). Indeed, as shown in **Figures 4** and **5**, for different input concentrations in the operational range the weight of the input can be tuned through the concentration of the enzyme DNA. To the best of our knowledge, the metabolic adders and perceptrons presented in this work are the first engineered biological circuits that use metabolism for analog computation.

Unlike genetic circuits that experience expression delays^2^, metabolic circuits have the advantage of faster response times since the enzymes have already been expressed in the system. Yet, metabolic circuits can be connected with the other layers of cellular information processing (like genetic or signal transduction layers) when needed, to build more complex sense-and-respond behaviors. The actuator layer of our perceptrons is a good example of this, where the calculated weighted sum is converted to fluorescence output via the genetic layer. In addition, we took advantage of the properties of cell-free systems, such as higher tunability and lack of toxicity^56, 64^, to rapidly build and characterize multiple combinations of transducer-actuator circuits. Cell-free systems can be lyophilized on paper and stored at ambient temperature for <1 year for diagnostic applications^16^. This expands the potential scope of cell-free metabolic perceptrons for use in multiplex detection of metabolic profiles in medical or environmental samples^16, 56^.

Here, we have built a single-layer perceptron, with positive weights, that can classify different profiles of input metabolites by applying different weights to each transducer. In the future, by adding competing or attenuating reactions that reduce the concentration of the transduced metabolite in response to an input, it may be possible to expand the training space by applying negative weights to certain inputs^65^. Furthermore, a single-layer perceptron can only classify data that is linearly separable^66^, which means that it should be possible to draw a line between the two classes of data points in order for the perceptron to classify them (**Supplementary Figure S7**). In contrast, multi-layer perceptrons, can approximate any function^67^ and can be used for more complex pattern recognition tasks^68^. With the use of bioretrosynthesis-based computational tools for metabolic pathway design, like Retropath^40^ and Sensipath^41^, it will be possible to build multiple layers of metabolic perceptrons that can classify complex patterns of metabolic states *in vivo*, or identify different metabolite concentrations in analytical samples. Finally, it may also be possible to apply *in situ* learning (within the whole-cell or cell-free environment) by applying winner selection strategies on successful classifiers^69^.

## Acknowledgments

AP is supported by INRA (French National Institute for Agricultural Research) and the idEx Paris-Saclay interdisciplinary doctoral fellowship. MKo is supported by DGA (French Ministry of Defense) and Ecole Polytechnique. PS is supported by ANR (grant number ANR-18-CE33-0015). JLF acknowledges support from BBSRC/EPSRC (grant number BB/M017702/1) and JLF and PS from the ANR (grant number ANR-18-CE33-0015).

## Author contributions

AP, MKo, MKu and JLF designed the project. AP designed and cloned the constructs, and performed the whole-cell experiments. AP, PLV, and JB designed cell-free experiment platform. AP, PLV, and PS performed cell-free experiments. MKo performed computational model simulations. All authors contributed to the manuscript write-up and approved the final manuscript.

## Methods

### Designing synthetic metabolic circuits

Retropath^40^ and Sensipath^41^ were used to design the metabolic circuits between potential input metabolites and detectable metabolites as outputs^47^. These tools function using a set of sink compounds, a set of source compounds, and a set of chemical rules^47, 70^ implementing enzyme-mediated chemical transformations. They then use retrosynthesis to propose pathways and the enzymes that can catalyze the necessary reactions, allowing promiscuity, between compounds from the sink and compounds from the source. To design the adder, the Retropath software was used with a set of detectable compounds as the sink and the molecules we wish to use as circuit inputs as the source. The results were potential pathways and the associated enzymes, which were then analyzed for feasibility. The sequences of the enzymes were codon-optimized, synthesized and implemented in *E. coli* or taken from a previous study.

### Molecular biology

All plasmids were made using Golden Gate assembly in *E. coli* Mach1 chemically competent cells. Whole-cell constructs were cloned in BioBrick standard vectors pSB1K3 (high-copy plasmid) and pSB4C5 (low-copy plasmid) and the TF and all the enzymes were constitutively expressed under constitutive promoter J23101 and RBS B0032. All cell-free plasmids were cloned in pBEAST^56^ (a derived vector from pBEST^71^). BenR cell-free plasmid and its cognate responsive prompter, pBen, expressing super-folder GFP were taken from our recent work^56^. All other cell-free enzymes were cloned under constitutive promoter J23101 and RBS B0032. Sequence and source of all the genes and parts are available in **Supplementary Table S5**. Synthetic sequences were provided by Twist Bioscience. Enzymes for cloning including Q5 DNA polymerase, BsaI, and T4 DNA ligase were purchased from New England Biolabs. DNA plasmids for cell-free reactions were prepared using the Macherey-Nagel maxiprep kit.

### Characterization of whole-cell circuits

For each circuit separate colonies of *E. coli* top10 strains harboring the circuit plasmids were cultured overnight at 37℃in LB with appropriate antibiotic. The next day each culture was diluted 100x in LB with antibiotics. 95 µL of fresh cultures were distributed in 96-well plate (Corning 3603) and the plate was incubated to reach the OD ∼ 0.1 in a plate reader (Biotek Synergy HTX). Then 5 µL of the input metabolites (100x ethanol solutions 5x diluted in LB) were added and the plate was incubated for 18 hours at 37℃. During the incubation, the OD_600_ and GFP fluorescence (gain: 35, ex: 458 nm, em: 528 nm) were measured. Benzoate, hippurate, cocaine hydrochloride, benzaldehyde, benzamide and biphenyl-2,3-diol (2,3-dihydroxy-biphenyl) were purchased from Sigma-Aldrich. Permission to purchase cocaine hydrochloride was given by the French drug regulatory agency (Agence Nationale de Sécurité du Médicament et des Produits de Santé). For all chemicals, serial dilutions of 100x concentrations were prepared in ethanol. The formula presenting the results of the circuits’ characterization is shown in data normalization section. The mean and standard deviation of all normalized data are provided in **Supplementary Table S6.**

### Cell-free extract and buffer preparation

Cell-free *E. coli* extract was produced as previously described^56, 72, 73^. Briefly, an overnight culture of BL21 Star (DE3)::RF1-CBD_3_ *E. coli* was used to inoculate 4L of 2xYT-P media in six 2 L flasks at a dilution of 1:100. The cultures were grown at 37°C with 220 rpm shaking for approximately 3.5-4 hours until the OD 600 = 2-3. Cultures were centrifuged at 5000 x g at 4°C for 12 minutes. Cell pellets were washed twice with 200 mL S30A buffer (14 mM Mg-glutamate, 60 mM K-glutamate, 50 mM Tris, pH 7.7), centrifuging after each wash at 5000 x g at 4°C for 12 minutes. Cell pellets were then resuspended in 40 mL S30A buffer and transferred to pre-weighed 50 mL Falcon conical tubes where they were centrifuged twice at 2000 x g at 4°C for 8 and 2 minutes, respectively, removing the supernatant after each. Finally, the tubes were reweighed and flash frozen in liquid nitrogen before storing at −80°C.

Cell pellets were thawed on ice and resuspended in 1 mL S30A buffer per gram of cell pellet. Cell suspensions were lysed via a single pass through a French press homogenizer (Avestin; Emulsiflex-C3) at 15000-20000 psi and then centrifuged at 12000 x g at 4°C for 30 minutes to separate out cellular cytoplasm. After centrifugation, the supernatant was collected and incubated at 37°C with 220 rpm shaking for 60 minutes. The extract was recentrifuged at 12000 x g at 4°C for 30 minutes, and the supernatant was transferred to 12-14 kDa MWCO dialysis tubing (Spectrum Labs; Spectra/Por4) and dialyzed against 2 L of S30B buffer (14 mM Mg-glutamate, 60 mM K-glutamate, ∼5 mM Tris, pH 8.2) overnight at 4°C. The following day, the extract was re-centrifuged one final time at 12000 x g at 4°C for 30 minutes, aliquoted, and flash frozen in liquid nitrogen before storage at −80°C.

The buffer for cell-free reactions is composed such that final reaction concentrations were as follows: 1.5 mM each amino acid except leucine, 1.25 mM leucine, 50 mM HEPES, 1.5 mM ATP and GTP, 0.9 mM CTP and UTP, 0.2 mg/mL tRNA, 0.26 mM CoA, 0.33 mM NAD, 0.75 mM cAMP, 0.068 mM folinic acid, 1 mM spermidine, 30 mM 3-PGA, and 2% PEG-8000. Additionally, the Mg-glutamate (0-6 mM), K-glutamate (20-140 mM), and DTT (0-3 mM) levels were serially calibrated for each batch of cell-extract for maximum signal. One batch of buffer was made for each batch of extract, aliquoted, and flash frozen in liquid nitrogen before storage at −80°C.

### Characterization of cell-free circuits

Cell-free reactions were performed in 15.75 µL of the mixture of 33.3% cell extract, 41.7% buffer, and 25% plasmid DNA, input metabolites, and water. The reactions were prepared in PCR tubes on ice and 15 µL of each was pipetted into 384-well plates (Thermo Scientific 242764). GFP fluorescence out of each circuit was recorded in the plate reader at 30℃ (gain: 50, ex: 458 nm, em: 528 nm). The background (cell-free reaction without any plasmid) corrected fluorescence data were normalized by 20 ng/µL of a plasmid expressing strong constitutive sfGFP (under OR2-OR1-Pr promoter^56^) and were plotted after 8 hours incubation. The mean and standard deviation of all normalized data are provided in **Supplementary Table S7.**

### Data normalization

For whole-cell data, we use the following normalization:

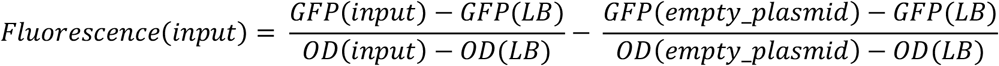

Reference: cells harboring empty plasmids

For cell-free data, we consider Relative Fluorescence Unit (RFU):

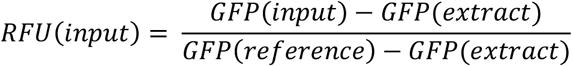

Reference: 20 ng/µL of a plasmid expressing the constitutive sfGFP under OR2-OR1-Pr promoter^56^.

### Simulation tools and parameter fitting

All data analysis and simulations were run on R (version 3.2.3)^74^. Dose-response curves were fitted using ordinary least squares errors and the R optim function (from Package stats version 3.2.3, using the L-BFGS-B method implementing the Limited-memory Broyden Fletcher Goldfarb Shanno algorithm, which is a quasi-Newton method). For the random parameter sampling around the mean fit, values were sampled from within +-1.96 standard error of the mean of the parameter estimation. The seed was set so as to ensure reproducibility. All simulations were run in the Rstudio development environment^75^. All parameters are presented in **Supplementary Tables S3** and **S4**.

### Whole-cell model

The whole-cell model is composed of three parts: the actuator, the transducers (which all obey the same law) and the resource competition.

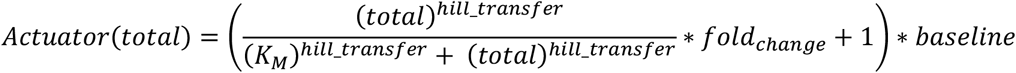

where *total* is the concentration of the considered input (in µM), *K_M_* is the concentration that allows for half-maximum induction (in µM), also termed IC_50_, *hill_transfer* is the Hill coefficient that characterizes the cooperativity of the induction system, fold_change is the dynamic range (in AU) and baseline is the basal GFP fluorescence without input (benzoate).

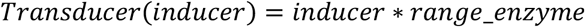

Where *input* is the input concentration in µM and *range_enzyme* is a dimensionless number characterizing the capacity of the enzyme to transduce the signal. When combining transducers with the actuator, transducer results are added before being fed into the actuator equation, just as benzoate concentrations are added before being converted to a fluorescent signal in the cell.

To account for resource competition, given our experimental results where there is little competition with one enzyme and significant competition with two, we used an equation including cooperativity of resource competition. This reduces the fold change of the actuator as there are less resources available for producing transcription factors and GFP.

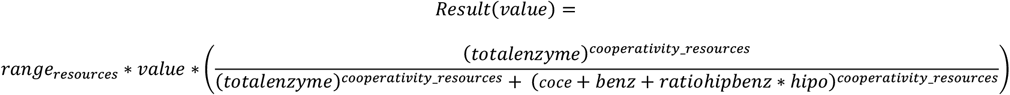

where *value* is the result of the actuator transfer function before accounting for resource competition, *range_resources, total_enzyme, cooperativity resources* characterize the Hill function that accounts for competition, *coce, benz and hipo* are the enzyme plasmid concentrations. *ratio_hip_benz* accounts for the differences in burden from different enzymes, its value around 0.8 is close to the ratio between enzyme lengths (1500 for benzaldehyde transducing enzyme and 1200 for HipO).

### Cell-free model

The model is composed of two parts: the actuator and the transducers.

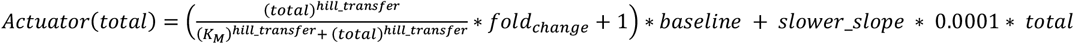

where *total* is the concentration of the considered input metabolite (in µM), *Km* is the concentration that allows for half-maximum induction (in µM), also termed IC50, *hill_transfer* is the Hill coefficient that characterizes the cooperativity of the induction system, *fold_change* is the dynamic range (in AU) and *baseline* is the basal GFP fluorescence without input (benzoate). *Slower_slope* accounts for the linearity observed in the actuator behavior at concentrations saturating the Hill transfer function.

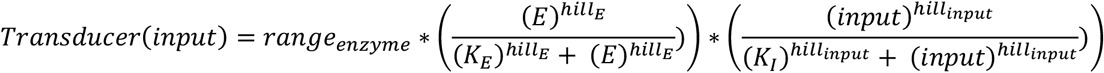

Where *range_enzyme* is a dimensionless number characterizing the capacity of the enzyme to transduce the signal. The activity of the enzyme is characterized by a Hill function as increasing concentrations do not lead to a linear increase but enzymes saturate (*E* is the enzyme quantity in nM, *KE* and *hillE* are its Hill constants), and similarly, *input* is the input metabolite concentration in µM with *KI* and *hill_input* as its Hill constants.

When combining transducers, transducer results are added before being fed into the actuator equation, just as benzoate concentrations are added before being converted to the fluorescent signal in the cell.

### Full model training process

Our training process is detailed in the Readme files supporting our modeling scripts provided in GitHub and is summarized here.

As the first step, the actuator transfer function model (benzoate transformed into fluorescence) is fitted 100 times on the actuator data, with all actuator parameters allowed to vary. The mean, standard deviation, standard error of the mean and confidence interval were saved at 95% of the estimation of those parameters. For transducer fitting (all transducers in cell-free and all except cocaine in whole-cell), we constrained the actuator characteristics in the following way: upper and lower allowed values are within the 95% confidence interval (or plus or minus one standard deviation from the mean for fold change and baseline in cell-free as it allowed a wider range, accounting for the decrease in actuator signal in transducer experiments without affecting the shape of the sigmoid). The initial values for the fitting process were sampled from a Gaussian distribution centered on the mean parameter estimation and spread with a standard deviation equal to the standard error of this parameter estimation. We then allowed fitting of all transducer parameters freely and of the actuator parameters within their 95% confidence interval.

Once this is done, all common parameters (actuator transfer function and resource competition) were sampled using the same procedure and fitting on the cocaine transducer was performed. To show that parameters are well constrained (proving they minimally explain the data), **Supplementary Figures S11** and **S12** show results of sampling parameters from the final parameters distribution (without fitting at that stage) and how they compare to the data.

### Objective functions and model scoring

In order to evaluate and compare our models, we used the following functions.

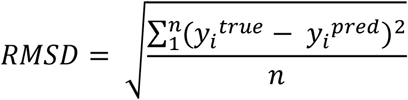

It measures how close the model is to the experiments. It allows for comparison of different models on the same data, the one with the smaller RMSD being better, but does not allow comparison between experiments.

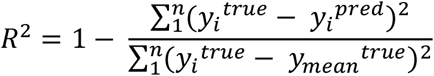

*R*^2^ allows measuring the goodness of fit. When the prediction is only around the sample mean, *R*^2^ = 0. When the predictions are close to the real experimental value, *R*^2^ gets closer to 1, whereas it can have important negative values when the model is really far off.

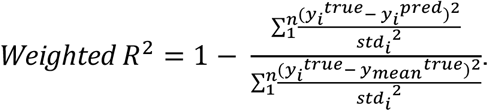

It is a variant of *R*^2^ that weights samples according to their experimental error, giving more weight o more certain samples. It otherwise has the same properties as *R*^2^.

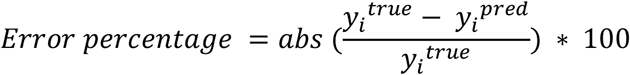

This measures the percentage of error for each point. We present the average on all experiments in Supplementary Tables S1 and S2.

### Perceptron weights calculation

In order to calculate the weights for the classifiers presented in **Figure 6**, we followed the following procedure. First, we defined the expected results (expressed in “OFF”s and “ON”s). We also defined a list of weights to test for each enzyme (here, between 0.1 nM and 10 nM, as tested in our weighted transducers). Then, for each combination of enzyme weights, we simulated the outcome of the classifiers for all possible input combinations. We then tested various possible thresholds and kept the enzyme combinations for which a threshold exists that allows for the expected behavior. As the last step, we manually analyzed the classifier to keep the ones both a high difference between ON and OFF, and a minimal enzyme weight to prevent resource competitions issues that could arise as we are adding more genes than previous experiments. In order to perform clusterings presented in **Supplementary Figure S8**, we sampled values uniformly within the stated ranges ([0, 2µM] for low values and [80, 100µM] for high values). We then simulated the results to assess the robustness of our designs.

### Binary clustering experiments

In order to perform the binary/2D clustering experiments, we sampled values uniformly within the stated ranges ([0, 2µM] for low values and [80, 100µM] for high values). For different weight (HipO and CocE) values, we simulated the fluorescence output of each of those cocaine-hippurate combinations. Moreover, for different threshold values (3, 3.5 and 4, as presented in **Supplementary Figure S7**), we numerically solved for the benzoate concentration such that

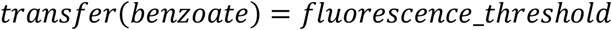

and then for values of cocaine and hippurate such that

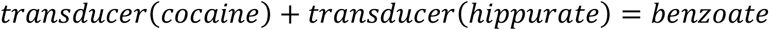

This equation with two unknowns gives us a curve of cocaine and hippurate values that would lie on our decided threshold for this set of weights. All combinations on the top right of that curve will be classified to “ON” and all combinations below will be classified to “OFF”.

### Code and data availability

All scripts and data for generating results presented in this paper are available at https://github.com/brsynth.

### Biological and chemical identifiers

In order to allow easier parsing of our article by bioinformatics tools, we provide here the identifiers of our biological sequences and chemical compounds.

*Benzoate (Benzoic acid):* InChI=1S/C7H6O2/c8-7(9)6-4-2-1-3-5-6/h1-5H,(H,8,9)

*Hippurate (Hippuric acid):* InChI=1S/C9H9NO3/c11-8(12)6-10-9(13)7-4-2-1-3-5-7/h1-5H,6H2,(H,10,13)(H,11,12)

*Cocaine:* InChI=1S/C17H21NO4/c1-18-12-8-9-13(18)15(17(20)21-2)14(10-12)22-16(19)11-6-4-3-5-7-11/h3-7,12-15H,8-10H2,1-2H3/t12-,13+,14-,15+/m0/s1

*Benzaldehyde:* InChI=1S/C7H6O/c8-6-7-4-2-1-3-5-7/h1-6H

*Biphenyl-2,3-diol:* InChI=1S/C12H10O2/c13-11-8-4-7-10(12(11)14)9-5-2-1-3-6-9/h1-8,13-14H

*Benzamide:* InChI=1S/C7H7NO/c8-7(9)6-4-2-1-3-5-6/h1-5H,(H2,8,9)

*BenR identifier:* UniProtKB - Q9L7Y6

*HipO identifier:* UniProtKB - P45493

*CocE identifier:* UniProtKB - Q9L9D7

*vdh identifier:* UniProtKB - D0RZT4

*bphC identifier:* UniProtKB - P17297

*bphD identifier:* UniProtKB - Q52036

*Benzamide transforming enzyme identifier:* UniProtKB - B4XEY3

Sequence and source of all the genes and parts are available in **Supplementary Table S5**

---

**Supplementary Figure S1.**
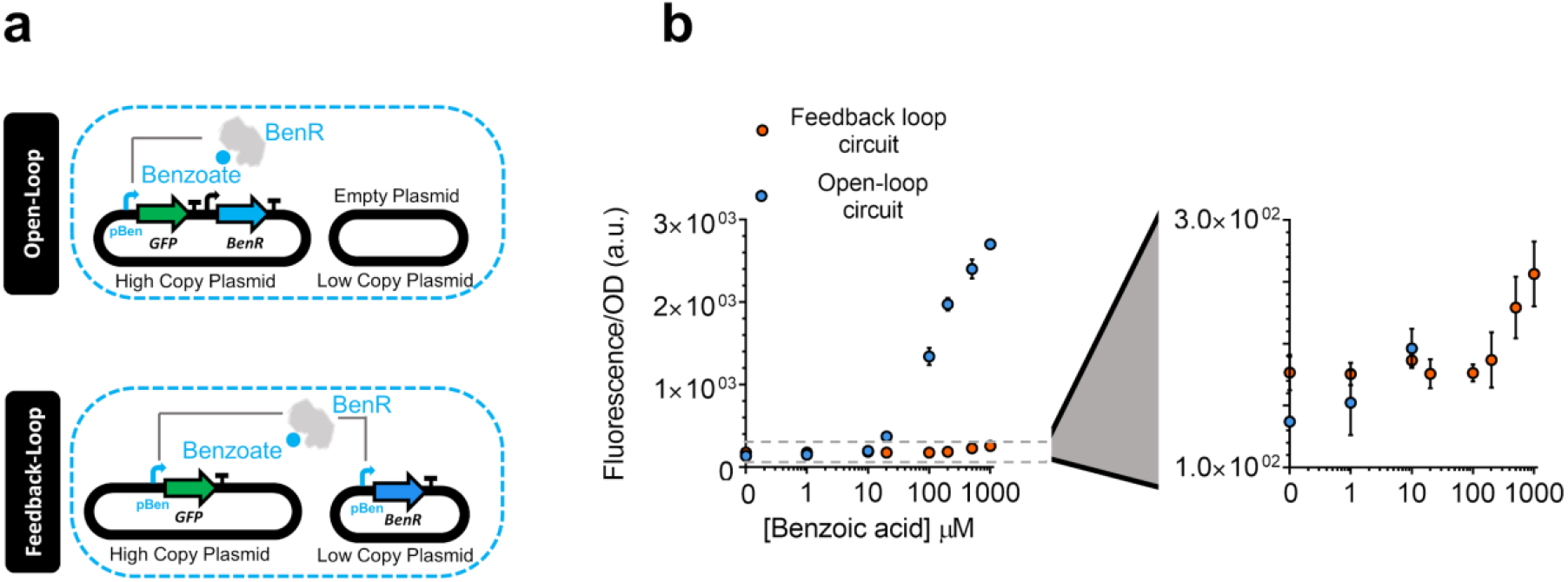
Feedback-loop circuit design of the benzoate actuator. (**a**) The open-loop circuit (Figure 1b) versus a feedback-loop circuit for the benzoate actuator. In the feedback-loop actuator the TF is expressed under its responsive promoter, pBen, in a low copy plasmid and sfGFP reporting the signal in a high copy plasmid1. (**b**) The dose-response of the feedback-loop versus the open-loop circuit (Figure 1c) to different concentrations of benzoate. All data points and the error bars are the mean and standard deviation of normalized values from three measurements.

**Supplementary Figure S2.**
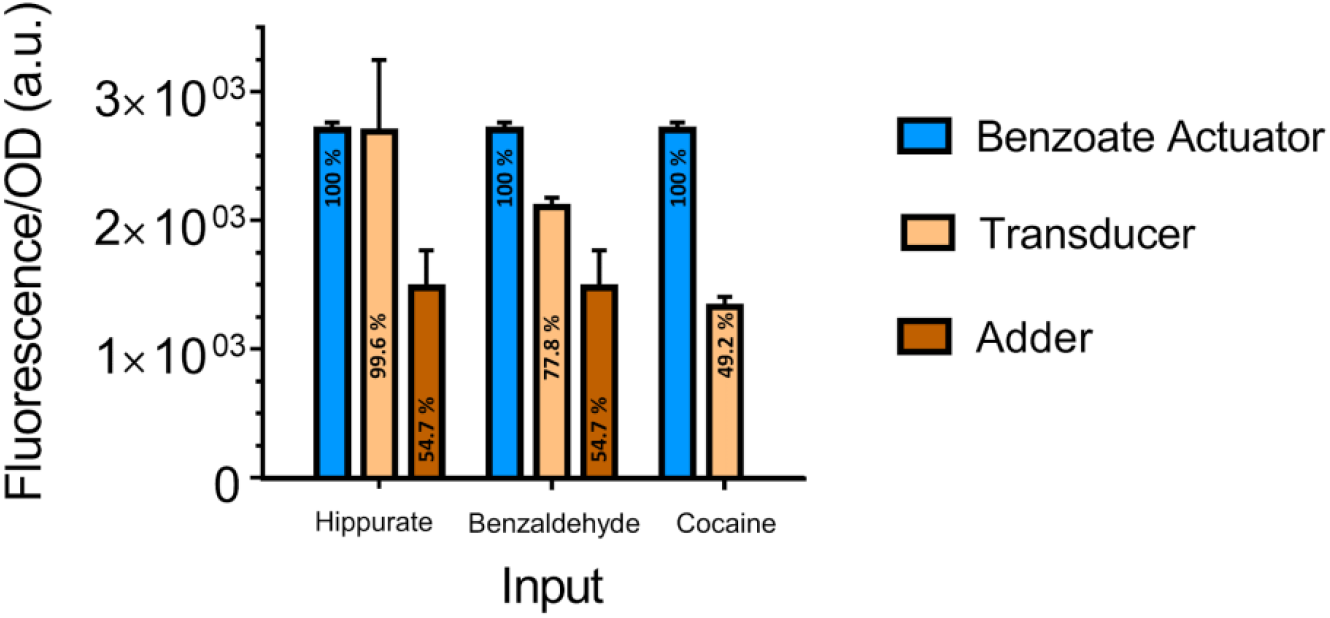
Comparison of the maximum signal of whole-cell circuits. Comparison of the maximal signal of hippurate, benzaldehyde, and cocaine transducers (beige) as well as hippurate-benzaldehyde adder (orange) with benzoate actuator (blue). The maximum signal of all the circuits are at maximum concentration of their inputs (1000 µM). The percentage in each bar represents its value with regard to maximum signal of benzoate in benzoate actuator. All data points and the error bars are from the results presented in Figures 1 and 2.

**Supplementary Figure S3.**
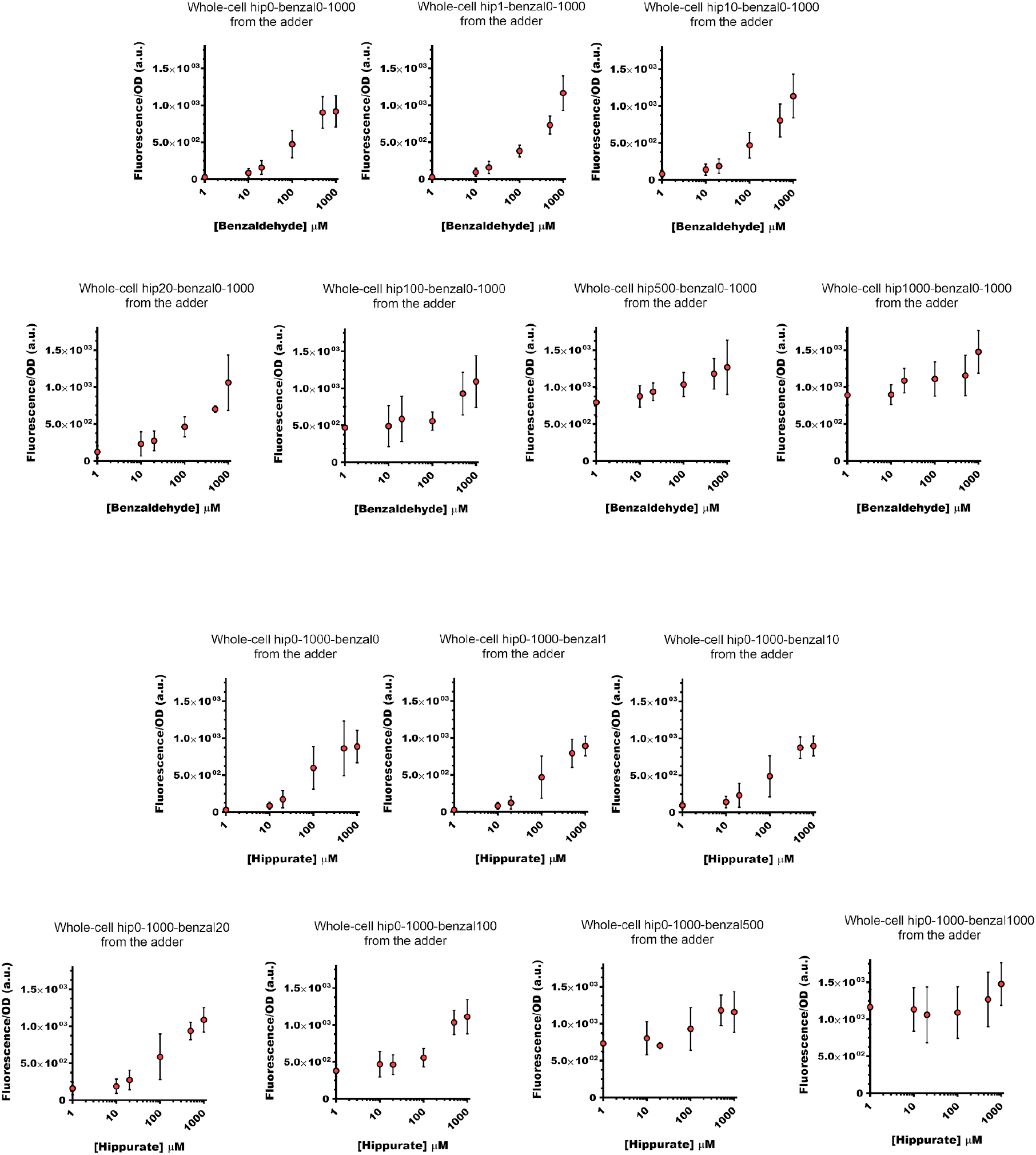
2D plots for the data presented in heatmap in Figure 2b. These 14 plots help visualize the linearity of metabolic addition. At the top of each plot the columns/rows of the heatmap in Figure 2b have been addressed.

**Supplementary Figure S4.**
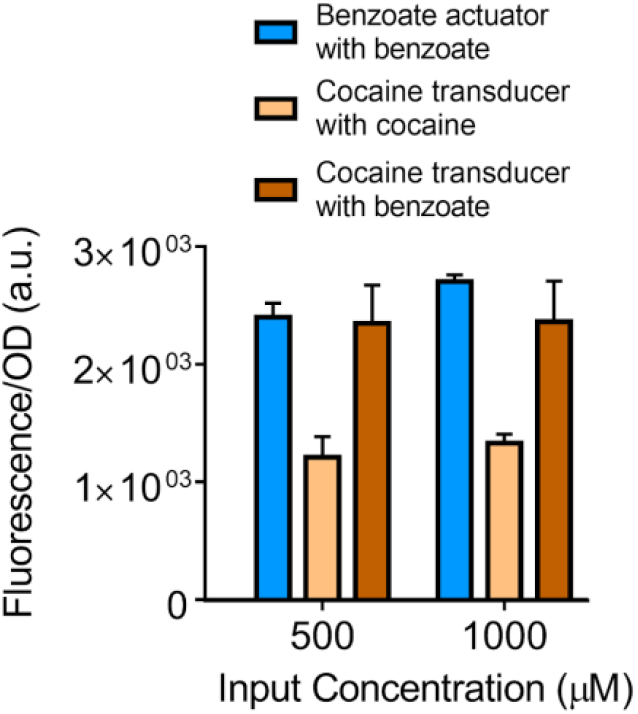
Examining the effect of resource competition on the whole-cell cocaine transducer. To study these effects on the single-enzyme metabolic circuit, the following experiment was performed: cocaine transducer (with the highest signal dissipation among the three tested in Figure 1) was supplied with benzoate input, to test the effect of enzymes on only cellular resource allocation but not conversion of inputs to benzoate. The cocaine transducer with benzoate input shows a behavior similar or close to the benzoate actuator. All data points and the error bars are the mean and standard deviation of normalized values from three measurements.

**Supplementary Figure S5.**
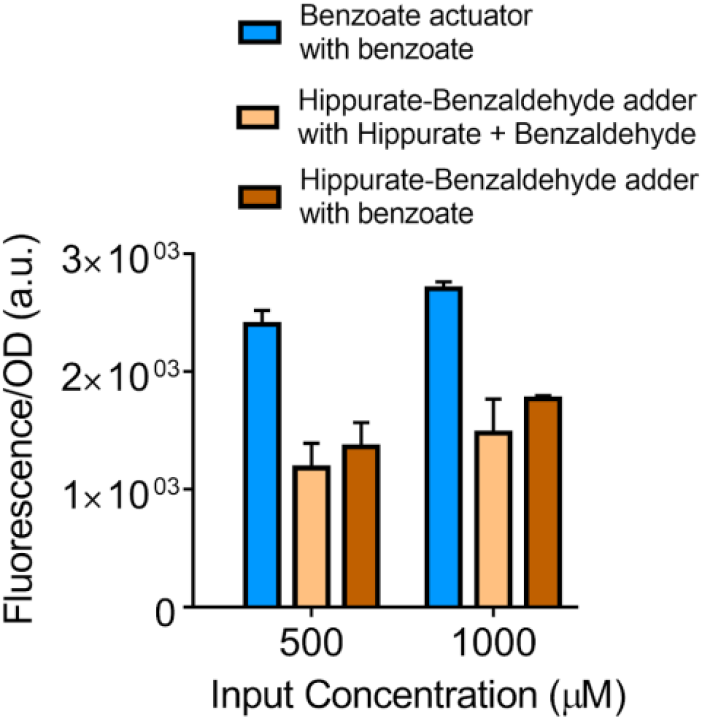
Examining the effect of enzyme efficiency on the whole-cell metabolic adder. To study these effects on the two-enzyme metabolic circuit (adder) the following experiment was performed: hippurate-benzaldehyde adder was supplied with benzoate input, to test the effect of enzymes on only cellular resource allocation but not conversion of inputs to benzoate. The adder with benzoate input shows a behavior similar to the adder inputted with hippurate and benzaldehyde. All data points and the error bars are the mean and standard deviation of normalized values from three measurements.

**Supplementary Figure S6.**
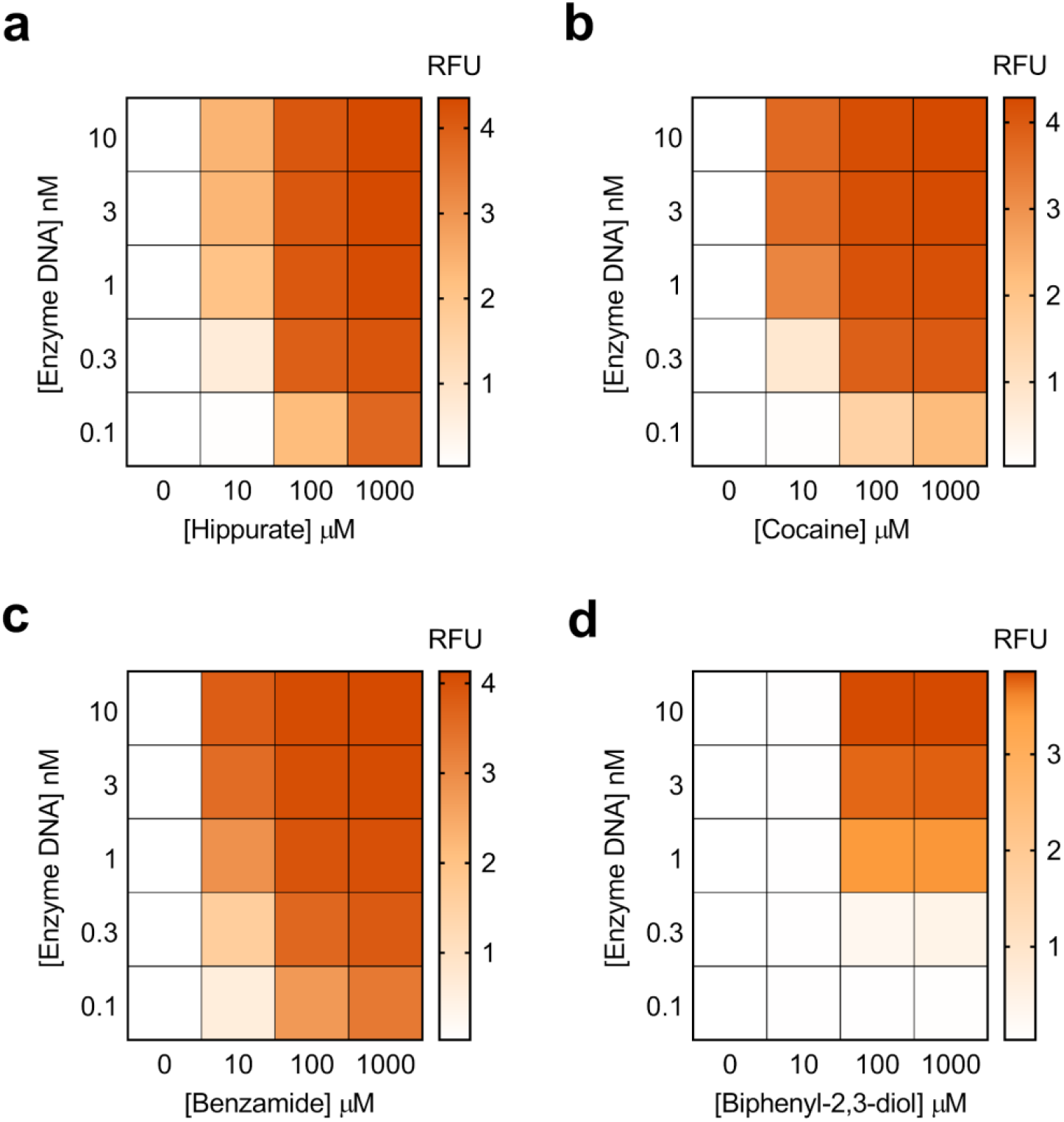
Weighted transducers model results. The model simulations for experimental conditions presented in Figure 4. (**a,b,c,d**) Heatmaps representing model simulations for weighted transducers at different concentrations of input molecules and enzymes DNA for hippurate (**a**), cocaine (**b**), benzamide (**c**) and biphenyl-2,3-diol (**d**).

**Supplementary Figure S7.**
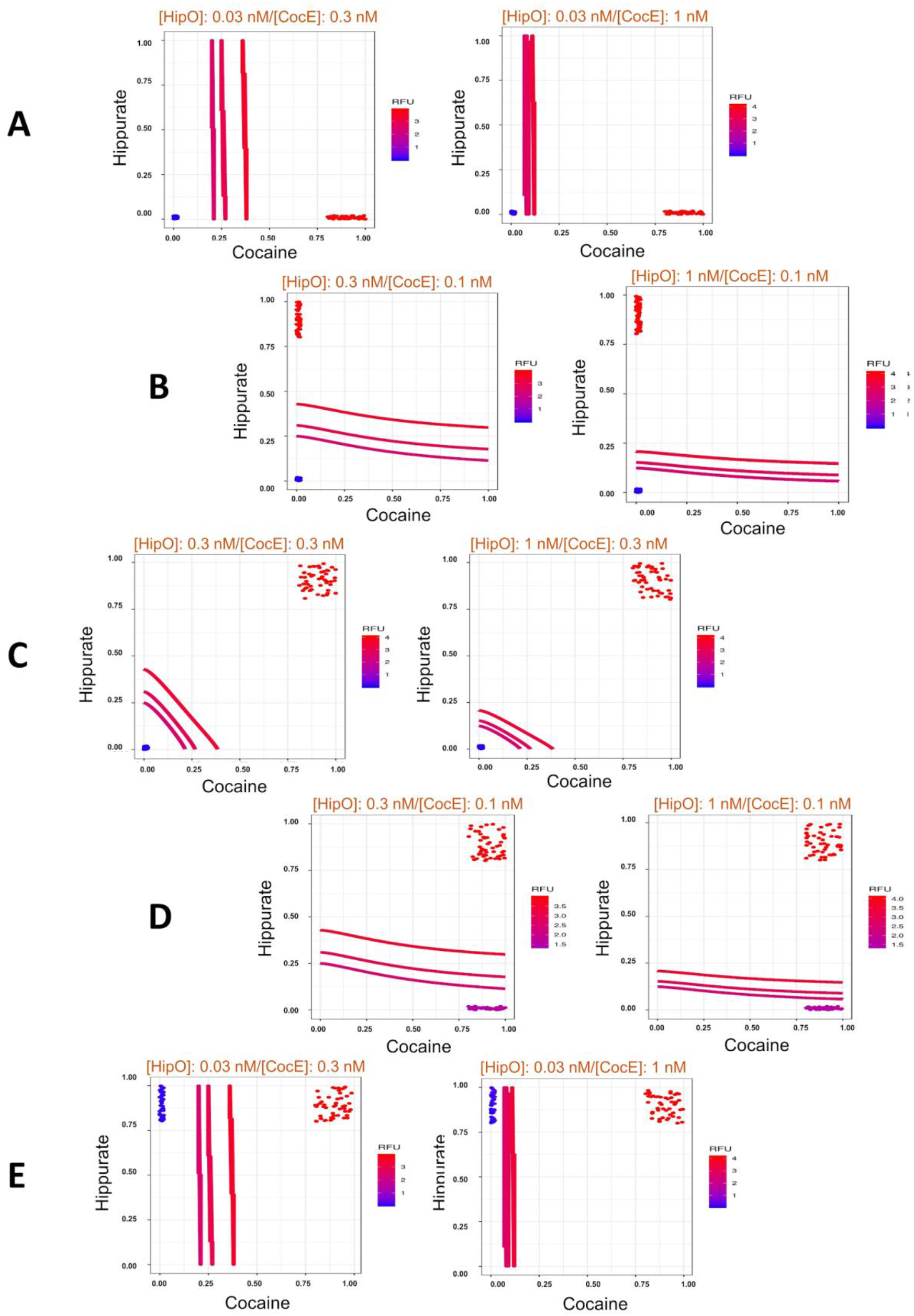
Five different binary classification problems using a metabolic perceptron for hippurate and cocaine. (**A** to **E**). For each problem, the scatter plot shows multiple data points that represent a combination of input values of cocaine and hippurate. The concentrations for those points are sampled between 0 and 2µM for low values and 80 and 100 µM for high values. The data points in each problem belong to two different sets that can be separated by a threshold line into two separate clusters. The trained model is then used to identify weights needed to be applied to the weighted transducers such that a decision threshold ‘d’ classifies the two clusters into red (ON, >d) or blue (OFF, <= d). The threshold lines shown in the plots represent three iso-fluorescence lines that successfully classify the data into the binary categories: ON and OFF.

**Supplementary Figure S8.**
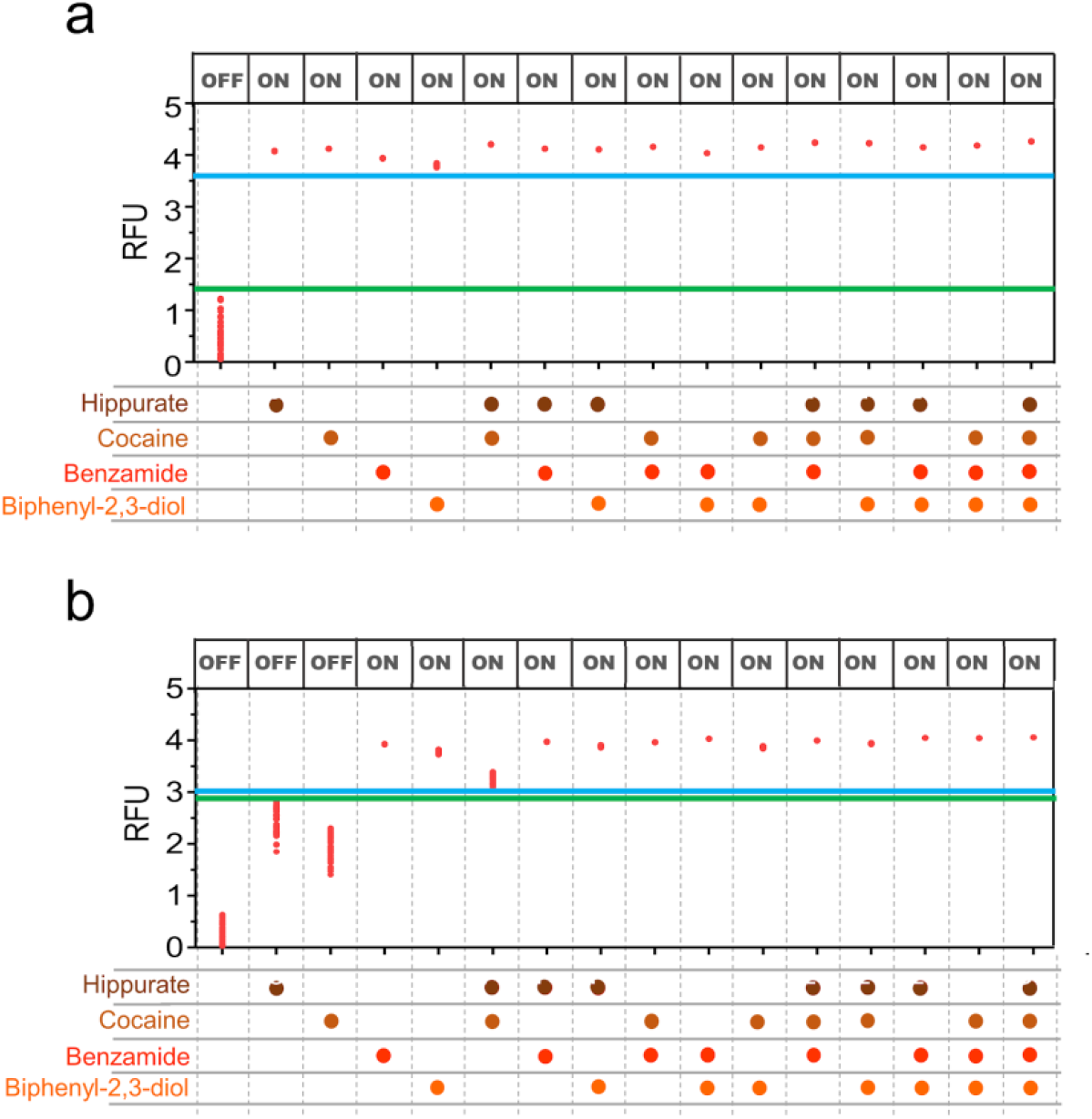
Model simulations for classifiers in Figure 6. Predictions associated with (**a**) the full OR classifier (Figure 6c) and (**b**) the first calculation for “[cocaine *(C)* AND hippurate *(H)*] OR benzamide *(B)* OR biphenyl-2,3-diol *(F)*” classifier with 0.1 nM HipO weight with (instead of 0.03 as experimentally tested and presented in Figure 6d). In order to perform the clusterings, we sampled values uniformly within the stated ranges ([0, 2µM] for low values and [80, 100µM] for high values). We then simulated the results to assess the robustness of our designs. The blue and green lines refer to the thresholds separating “OFF” and “ON” states. The panel of “OFF” and “ON” at the top of the plots are the expected outputs. (RFU: Relative Fluorescence Unit).

**Supplementary Figure S9.**
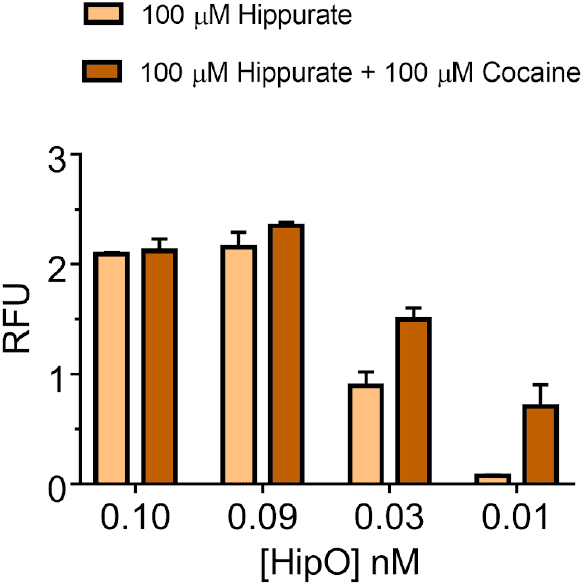
Further characterization of HipO enzyme (hippurate transforming enzyme) at lower concentrations of the enzyme and 100 µM hippurate. HipO enzyme which for its weight led to higher signals than predicted, needed to be further characterized at concentrations lower than the minimum concentration used for the weighted metabolic circuits (0.1 nM). For this characterization, this figure shows the effect of 100 µM hippurate input alone and its additive effect when coupled with 100 µM cocaine at the weight (CocE enzyme concentration) of 0.1 nM. All data are the mean and the error bars are the standard deviation of normalized values from three measurements. (RFU: Relative Fluorescence Unit).

**Supplementary Figure S10.**
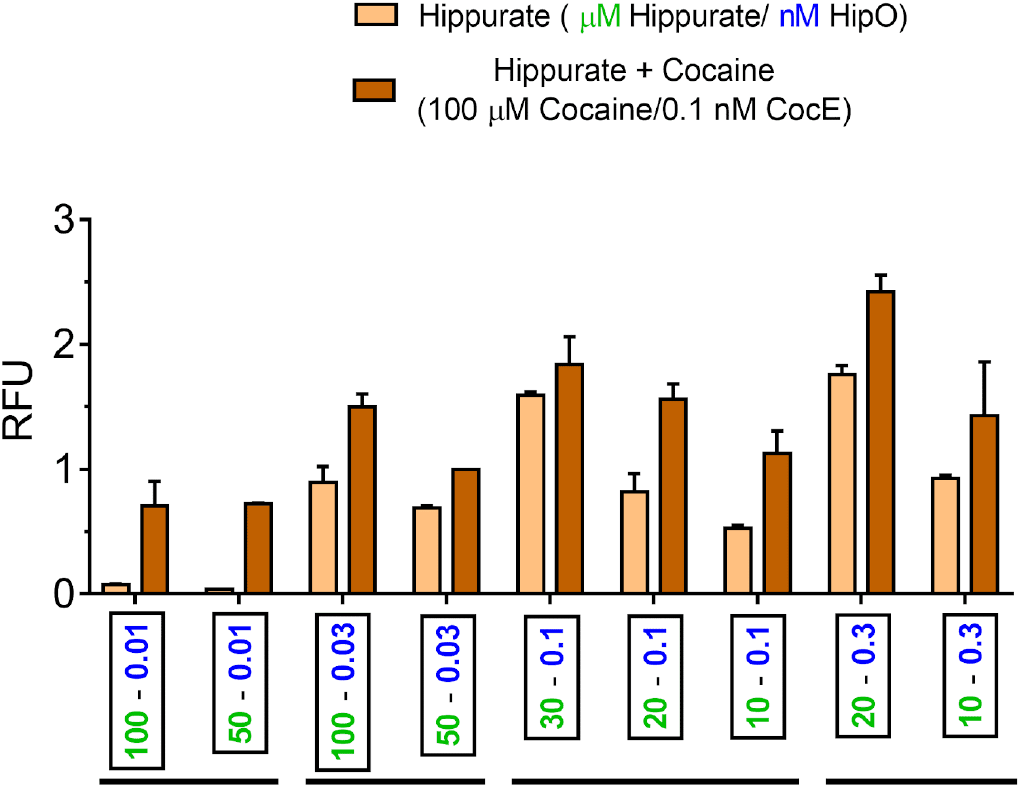
Exploring Hippurate-Cocaine ON-OFF behavior with different weights and input concentrations for hippurate. All these experiments were done while Cocaine is at concentration of 100 uM and weight of 0.1 nM CocE. All data are the mean and the error bars are the standard deviation of normalized values from three measurements. (RFU: Relative Fluorescence Unit).

**Supplementary Figure S11.**
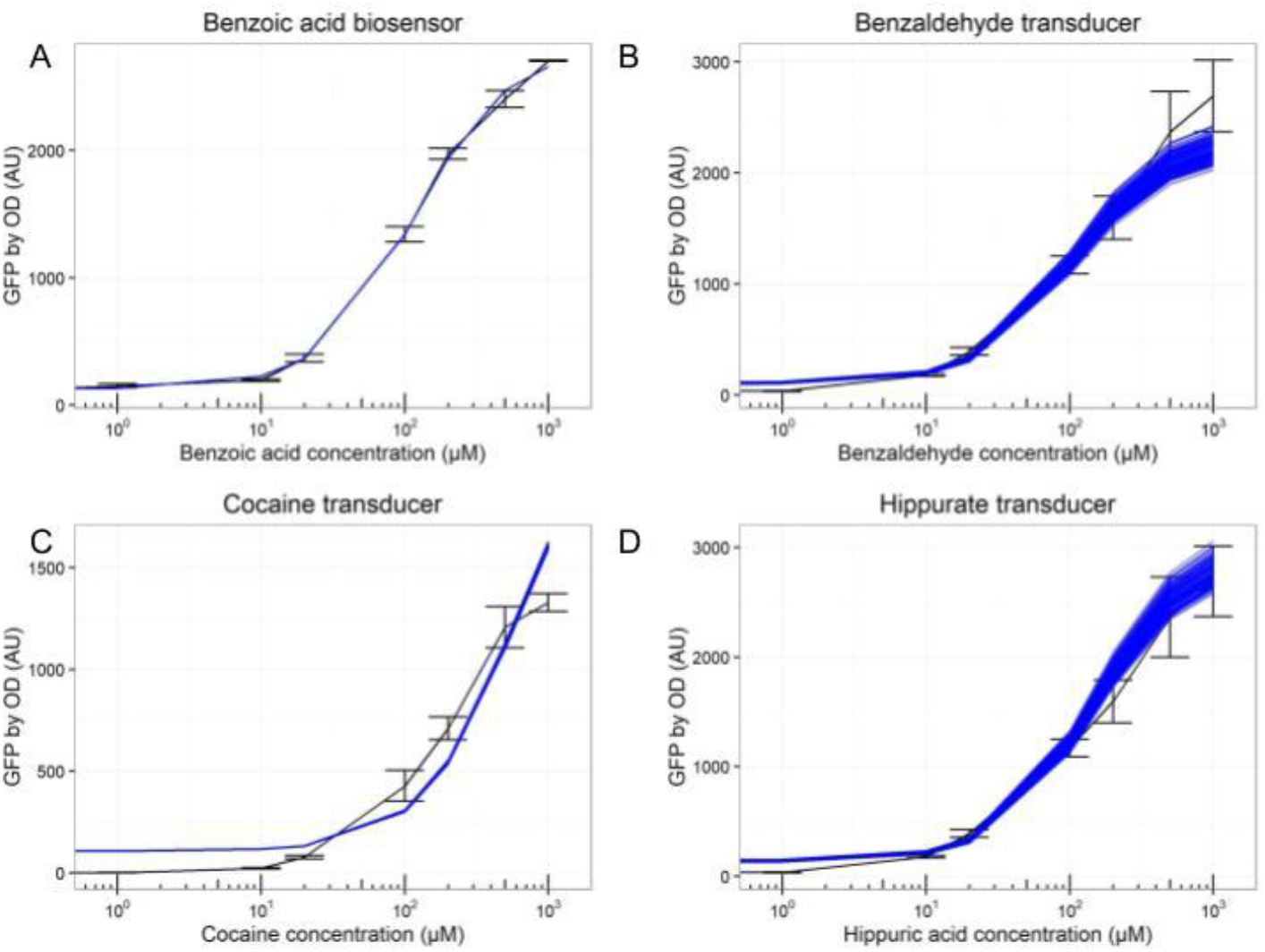
Simulations from the random sampling of estimated parameters in whole-cell system. Representation of the experimental data with SEM (n = 3) in black, and in blue, the results from 100 simulations of the model with parameters drawn from the final parameters estimation without refitting. The combination of various parameters within our estimations correctly recapitulates the data. (A) benzoate actuator, (B) benzaldehyde transducer, *(B)* cocaine transducer, and (D) hippurate transducer. Scripts provided in GitHub also allow for visualization of those results for each axis of the adder in Figure 2.

**Supplementary Figure S12.**
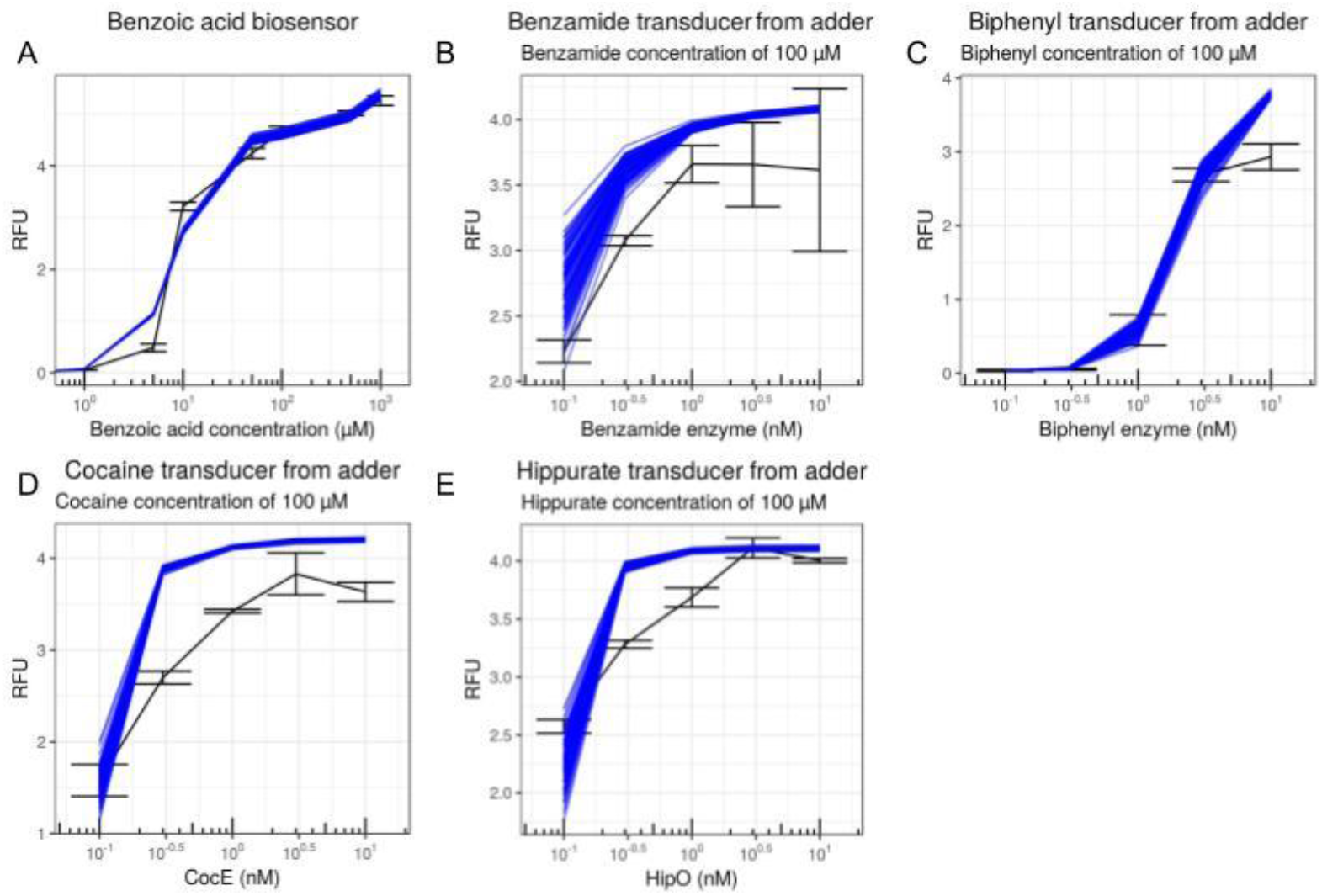
Simulations from the random sampling of estimated parameters in the cell-free system. Representation of the experimental data with SEM (n = 3) in black, and in blue, the results from 100 simulations of the model with parameters drawn from the final parameters estimation without refitting. The combination of various parameters within our estimations correctly recapitulates the data. (A) benzoate actuator, (B) benzamide transducer, (C) biphenyl-2,3-diol transducer, (D) cocaine transducer, and (E) hippurate transducer. The simulation of the transducers were performed with 100 µM of the input metabolites as will be used in the classifier experiments. Scripts provided in GitHub also allow for visualisation of those results for other axis of the various heatmaps in Figure 4. (RFU: Relative Fluorescent/expression Unit of GFP).

**Supplementary Table S1.**
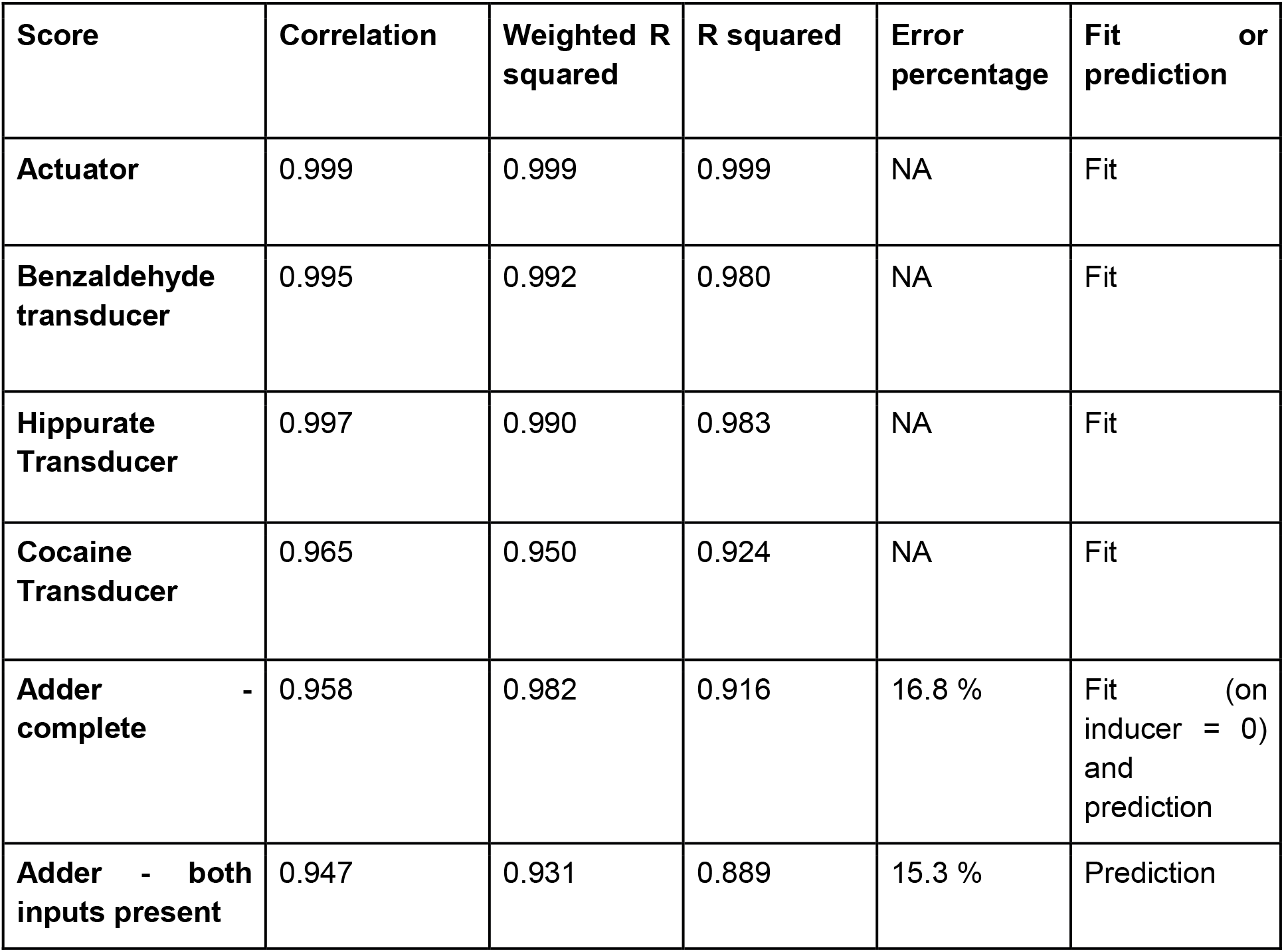
Goodness of fit scores for the whole-cell models. The correlation (from the R cor function), Weighted R squared and R squared between the experimental data and the model. Exact definition of the weighted R squared and the R squared are provided in the Methods section, as well as the RMSD that is used to compare models.

**Supplementary Table S2.**
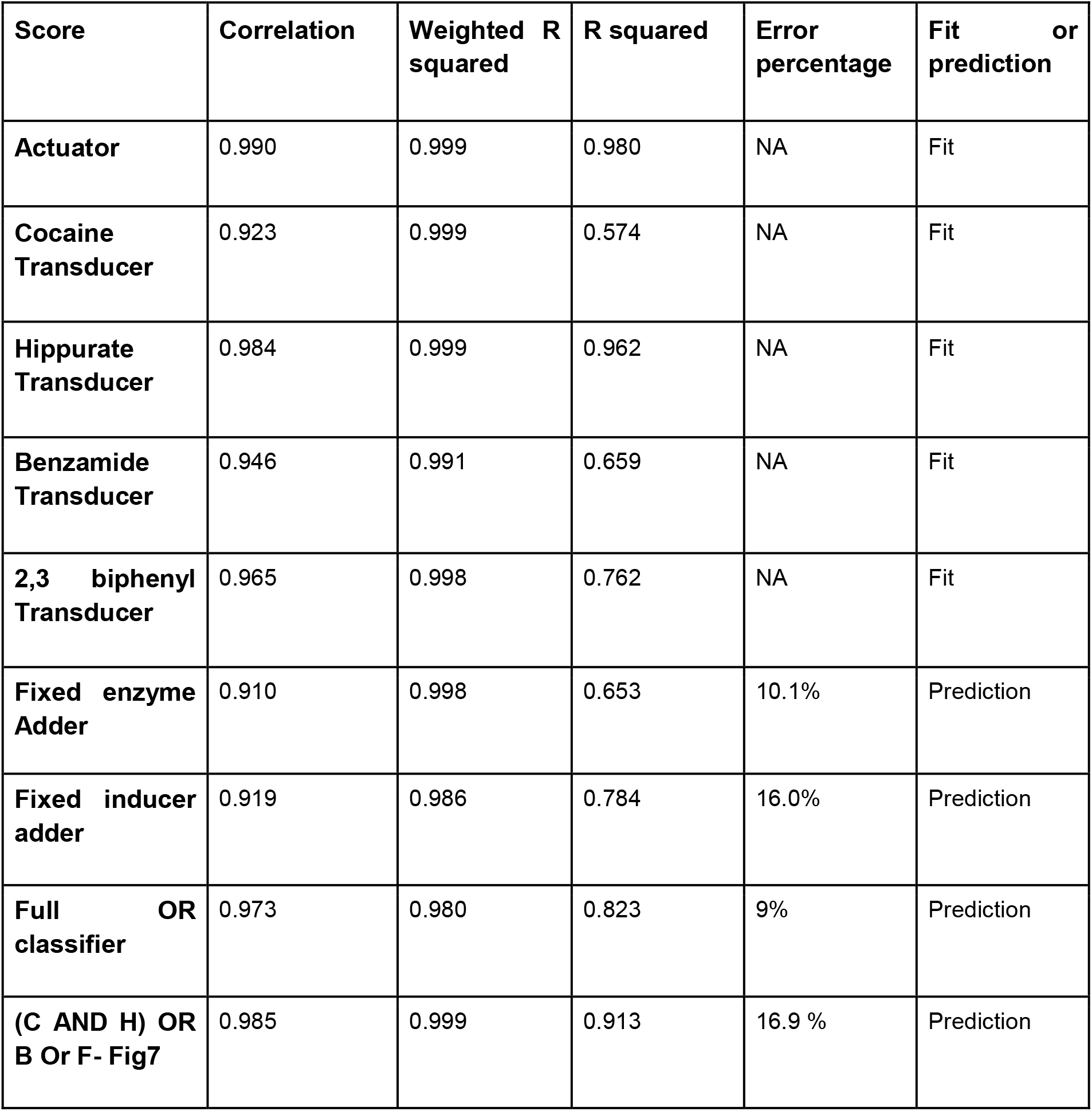
Goodness of fit scores for the cell-free models.

| **Score** | **Correlation** | **Weighted R squared** | **R squared** | **Error percentage** | **Fit or prediction** |
| --- | --- | --- | --- | --- | --- |
| **Actuator** | 0.990 | 0.999 | 0.980 | NA | Fit |
| **Cocaine Transducer** | 0.923 | 0.999 | 0.574 | NA | Fit |
| **Hippurate Transducer** | 0.984 | 0.999 | 0.962 | NA | Fit |
| **Benzamide Transducer** | 0.946 | 0.991 | 0.659 | NA | Fit |
| **2,3 biphenyl Transducer** | 0.965 | 0.998 | 0.762 | NA | Fit |
| **Fixed enzyme Adder** | 0.910 | 0.998 | 0.653 | 10.1% | Prediction |
| **Fixed inducer adder** | 0.919 | 0.986 | 0.784 | 16.0% | Prediction |
| **Full OR classifier** | 0.973 | 0.980 | 0.823 | 9% | Prediction |
| **(C AND H) OR B Or F- Fig7** | 0.985 | 0.999 | 0.913 | 16.9 % | Prediction |

**Supplementary Table S3.** Parameter estimations for in vivo model. Mean value plus and minus 95% Confidence Interval

| Parameter | Mean Value +- 95 Confidence Interval |
| --- | --- |
| Hill_transfer | 1.34 +- 1 e-6 |
| Km | 114 +- 1 e-4 |
| Fold_change | 20.6 +- 3 e-5 |
| Baseline | 130 +- 2 e-4 |
| Range_BenZ | 1.1 +- 1 e-6 |
| Range_HipO | 0.787 +- 1 e-6 |
| Range_CocE | 0.201 +- 2.97 e-3 |
| total_enzyme | 4.22 +- 0.193 |
| Ratio_hip_benz | 0.776 +- 3.7 e-3 |
| Cooperativity_resource | 1.956 +- 4.56 e-2 |
| Range_resource | 1.973 +- 0.107 |

**Supplementary Table S4.** Parameter estimations for cell-free model. Mean value plus and minus 95% Confidence Interval (Standard Deviation for fold change and baseline)

| Parameter | Mean Value +- 95 CI |
| --- | --- |
| Hill_transfer | 2.2 +- 0.1 |
| Km | 8.40 +- 9 e-3 |
| Fold_change | 137 +- 1.84 (sd : 9.41) |
| Baseline | 3.29 e-2 +- 4 e-4 (sd : 2 e-3 ) |
| Slower_slope | 8.19 +- 9.3 e-2 |
| Range_HipO | 488 +- 35 |
| HipO_constant | 0.396 +- 0.022 |
| Hippurate_constant | 245 +- 29 |
| Hill_HipO | 1.82 +- 0.052 |
| Hill_hippurate | 1.205 +- 0.046 |
| Range_CocE | 337 +- 28 |
| CocE_constant | 0.799 +- 0.00017 |
| Cocaine_constant | 54 .4 +- 5.04 |
| Hill_CocE | 1.713 +- 0.055 |
| Hill_cocaine | 1.44 +- 0.047 |
| range_benzamid_enz | 234 +- 20 |
| benzamid_enz_constant | 3.73 +- 0.27 |
| benzamid_constant | 48.6 +- 5.5 |
| hill_benzamid_enz | 0.683 +- 0.072 |
| hill_benzamid | 0.906 +- 0.087 |
| range_biphenyl_enz | 63.7 +- 4.79 |
| biphenyl_enz_constant | 8.63 +- 0.31 |
| biphenyl_constant | 56.3 +- 4.92 |
| hill_biphenyl_enz | 1.25 +- 0.067 |
| hill_biphenyl | 3.05 +- 0.192 |

**Supplementary Table S5.**
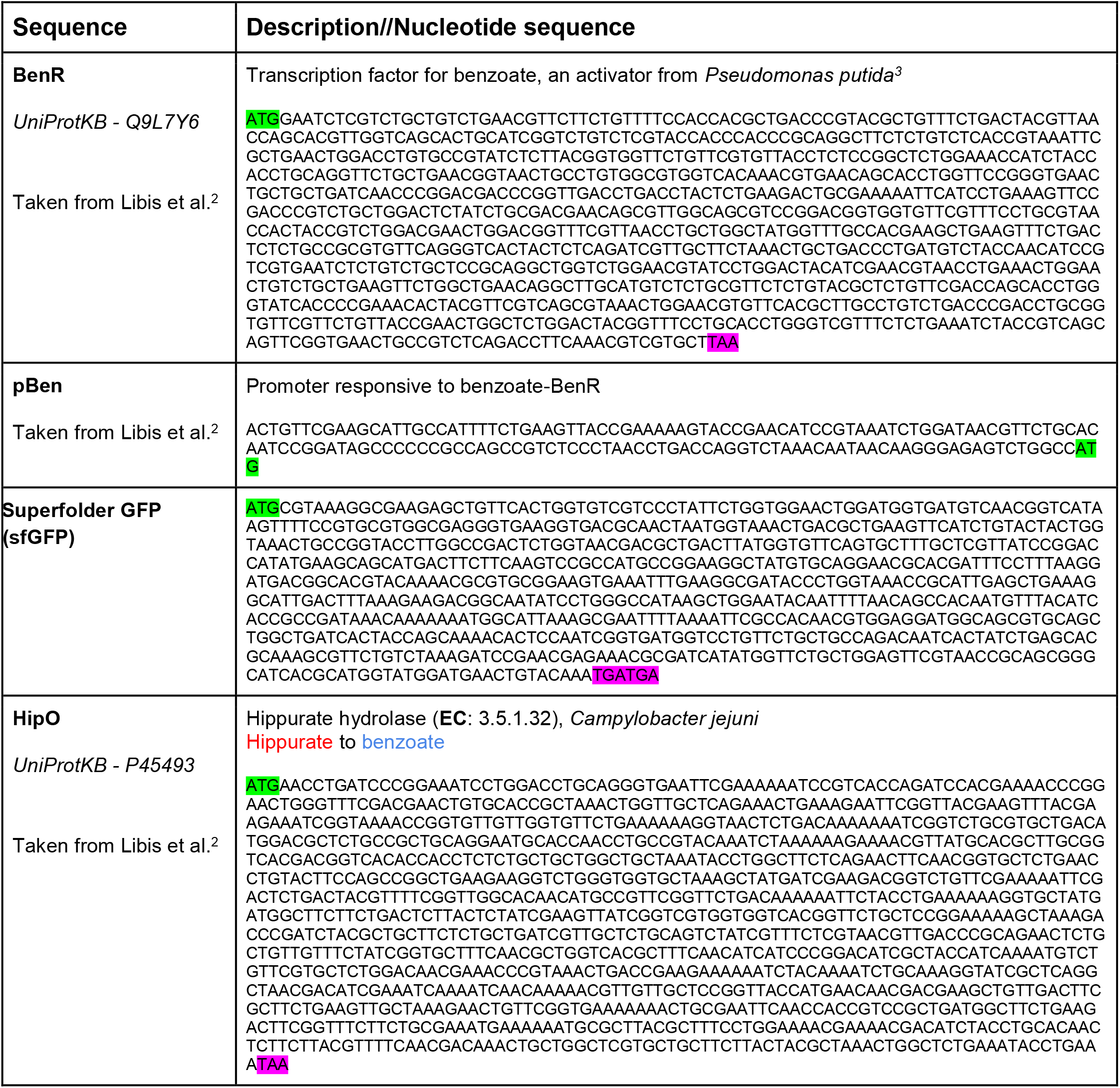
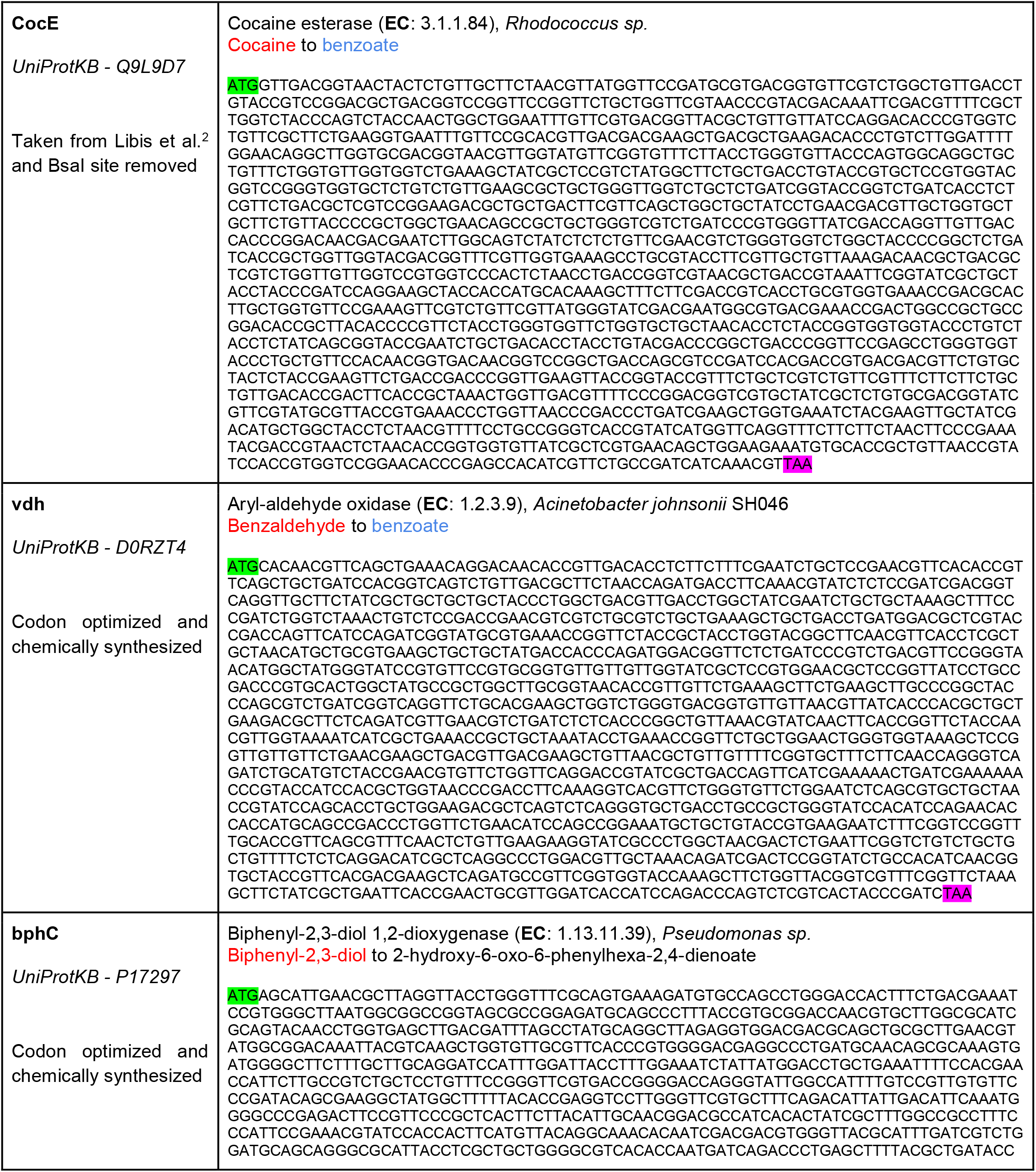
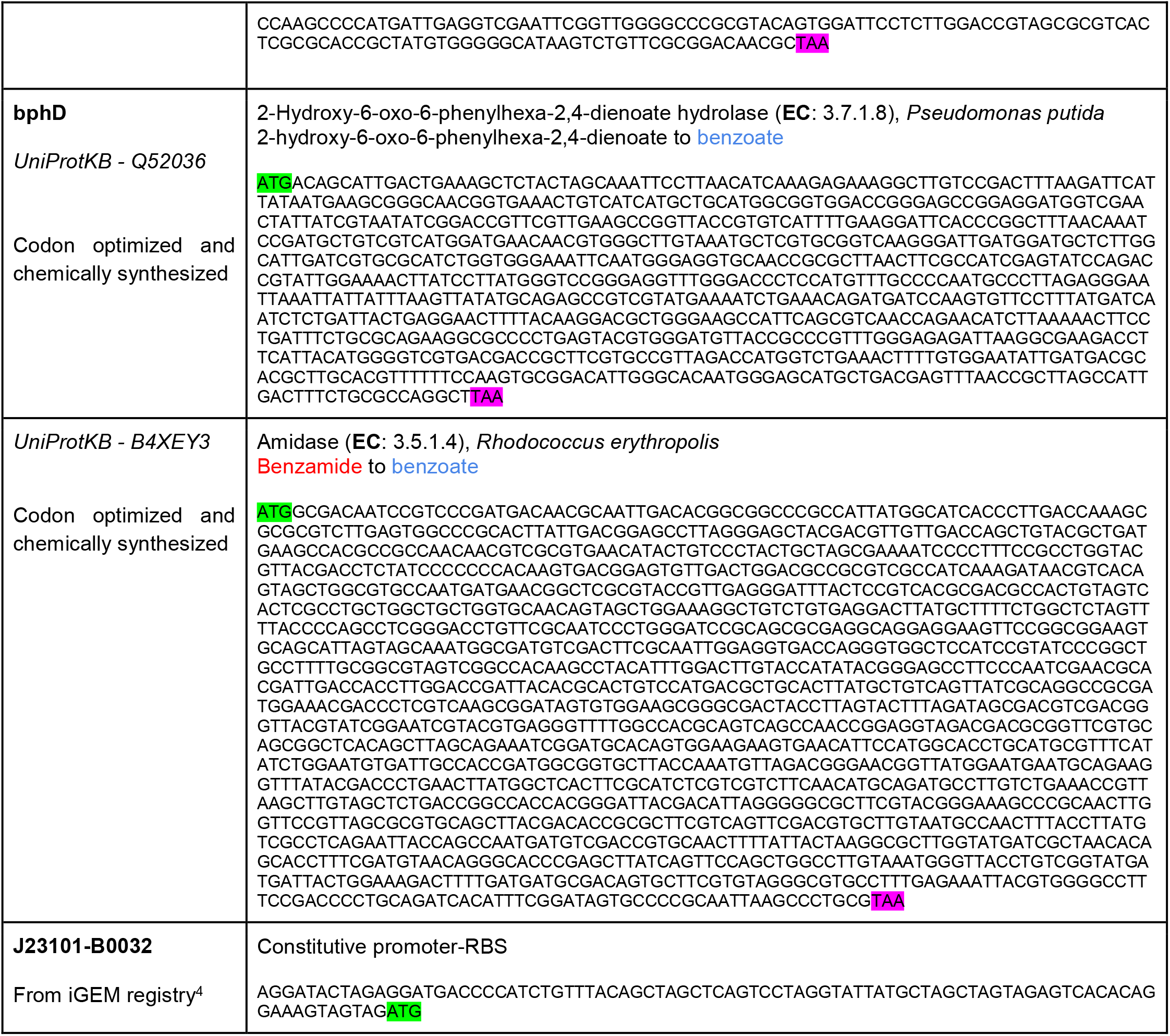
List of sequences and their source used in this study.

**Supplementary Table S6**. Excel file containing the mean and standard deviation of the normalized data of <u>whole-cell experiments</u>, and model simulated/predicted results associated with each experiment.

**Supplementary Table S7**. Excel file containing the mean and standard deviation of the normalized data of <u>cell-free experiments</u> and model simulated/predicted results associated with each experiment.

## References

1. Purcell, O. & Lu, T. K. Synthetic analog and digital circuits for cellular computation and memory. Curr. Opin. Biotechnol. 29, 146–155 (2014).

2. Brophy, J. A. N. & Voigt, C. A. Principles of genetic circuit design. Nat. Methods 11, 508–520 (2014).

3. Purnick, P. E. M. & Weiss, R. The second wave of synthetic biology: from modules to systems. Nat. Rev. Mol. Cell Biol. 10, 410–422 (2009).

4. Selberg, J., Gomez, M. & Rolandi, M. The Potential for Convergence between Synthetic Biology and Bioelectronics. Cell Syst 7, 231–244 (2018).

5. Moon, T. S., Lou, C., Tamsir, A., Stanton, B. C. & Voigt, C. A. Genetic programs constructed from layered logic gates in single cells. Nature 491, 249–253 (2012).

6. Shis, D. L., Hussain, F., Meinhardt, S., Swint-Kruse, L. & Bennett, M. R. Modular, Multi-Input Transcriptional Logic Gating with Orthogonal LacI/GalR Family Chimeras. ACS Synthetic Biology 3, 645–651 (2014).

7. Buffi, N. et al. Miniaturized bacterial biosensor system for arsenic detection holds great promise for making integrated measurement device. Bioeng. Bugs 2, 296–298 (2011).

8. Wan, X. et al. Cascaded amplifying circuits enable ultrasensitive cellular sensors for toxic metals. Nat. Chem. Biol. (2019). doi:10.1038/s41589-019-0244-3

9. Courbet, A., Endy, D., Renard, E., Molina, F. & Bonnet, J. Detection of pathological biomarkers in human clinical samples via amplifying genetic switches and logic gates. Sci. Transl. Med. 7, 289ra83 (2015).

10. Wen, K. Y. et al. A Cell-Free Biosensor for Detecting Quorum Sensing Molecules in P. aeruginosa-Infected Respiratory Samples. ACS Synthetic Biology 6, 2293–2301 (2017).

11. Kemmer, C. et al. Self-sufficient control of urate homeostasis in mice by a synthetic circuit. Nat. Biotechnol. 28, 355–360 (2010).

12. Isabella, V. M. et al. Development of a synthetic live bacterial therapeutic for the human metabolic disease phenylketonuria. Nat. Biotechnol. 36, 857–864 (2018).

13. Zhang, F., Carothers, J. M. & Keasling, J. D. Design of a dynamic sensor-regulator system for production of chemicals and fuels derived from fatty acids. Nat. Biotechnol. 30, 354–359 (2012).

14. Koch, M., Pandi, A., Borkowski, O., Batista, A. C. & Faulon, J.-L. Custom-made transcriptional biosensors for metabolic engineering. Current Opinion in Biotechnology 59, 78–84 (2019).

15. Nielsen, A. A. K. et al. Genetic circuit design automation. Science 352, aac7341 (2016).

16. Pardee, K. et al. Paper-Based Synthetic Gene Networks. Cell 159, 940–954 (2014).

17. Courbet, A., Amar, P., Fages, F., Renard, E. & Molina, F. Computer-aided biochemical programming of synthetic microreactors as diagnostic devices. Mol. Syst. Biol. 14, e8441 (2018).

18. Katz, E. Enzyme-Based Logic Gates and Networks with Output Signals Analyzed by Various Methods. Chemphyschem 18, 1688–1713 (2017).

19. Kiel, C., Yus, E. & Serrano, L. Engineering Signal Transduction Pathways. Cell 140, 33–47 (2010).

20. Shaw, W. M. et al. Engineering a Model Cell for Rational Tuning of GPCR Signaling. Cell (2019). doi:10.1016/j.cell.2019.02.023

21. Daniel, R., Rubens, J. R., Sarpeshkar, R. & Lu, T. K. Synthetic analog computation in living cells. Nature 497, 619–623 (2013).

22. Farzadfard, F. & Lu, T. K. Synthetic biology. Genomically encoded analog memory with precise in vivo DNA writing in living cell populations. Science 346, 1256272 (2014).

23. Bonnet, J., Subsoontorn, P. & Endy, D. Rewritable digital data storage in live cells via engineered control of recombination directionality. Proc. Natl. Acad. Sci. U. S. A. 109, 8884–8889 (2012).

24. Bonnet, J., Yin, P., Ortiz, M. E., Subsoontorn, P. & Endy, D. Amplifying genetic logic gates. Science 340, 599–603 (2013).

25. Zeng, J. et al. A Synthetic Microbial Operational Amplifier. ACS Synth. Biol. 7, 2007–2013 (2018).

26. Green, A. A., Silver, P. A., Collins, J. J. & Yin, P. Toehold Switches: De-Novo-Designed Regulators of Gene Expression. Cell 159, 925–939 (2014).

27. Bikard, D. et al. Programmable repression and activation of bacterial gene expression using an engineered CRISPR-Cas system. Nucleic Acids Res. 41, 7429–7437 (2013).

28. Nielsen, A. A. K. & Voigt, C. A. Multi-input CRISPR/Cas genetic circuits that interface host regulatory networks. Mol. Syst. Biol. 10, 763 (2014).

29. Silva-Rocha, R., Tamames, J., dos Santos, V. M. & de Lorenzo, V. The logicome of environmental bacteria: merging catabolic and regulatory events with Boolean formalisms. Environ. Microbiol. 13, 2389–2402 (2011).

30. Goñi-Moreno, A. & Nikel, P. I. High-Performance Biocomputing in Synthetic Biology– Integrated Transcriptional and Metabolic Circuits. Front. Bioeng. Biotechnol. 7, (2019).

31. Prindle, A. et al. Rapid and tunable post-translational coupling of genetic circuits. Nature 508, 387–391 (2014).

32. Cheng, Y.-Y., Hirning, A. J., Josić, K. & Bennett, M. R. The Timing of Transcriptional Regulation in Synthetic Gene Circuits. ACS Synth. Biol. 6, 1996–2002 (2017).

33. Valdez-Cruz, N. A., Caspeta, L., Pérez, N. O., Ramírez, O. T. & Trujillo-Roldán, M. A. Production of recombinant proteins in E. coli by the heat inducible expression system based on the phage lambda pL and/or pR promoters. Microbial Cell Factories 9, 18 (2010).

34. Anderson, J. C., Clarke, E. J., Arkin, A. P. & Voigt, C. A. Environmentally controlled invasion of cancer cells by engineered bacteria. J. Mol. Biol. 355, 619–627 (2006).

35. Tabor, J. J. et al. A synthetic genetic edge detection program. Cell 137, 1272–1281 (2009).

36. Meyer, A. J., Segall-Shapiro, T. H., Glassey, E., Zhang, J. & Voigt, C. A. Escherichia coli ‘Marionette’ strains with 12 highly optimized small-molecule sensors. Nature Chemical Biology 15, 196–204 (2019).

37. Sauro, H. M. & Kim, K. H. Synthetic biology: It’s an analog world. Nature 497, 572–573 (2013).

38. Daniel, R., Woo, S. S., Turicchia, L. & Sarpeshkar, R. Analog transistor models of bacterial genetic circuits. 2011 IEEE Biomedical Circuits and Systems Conference (BioCAS) (2011). doi:10.1109/biocas.2011.6107795

39. Rosenblatt, F. The perceptron: a probabilistic model for information storage and organization in the brain. Psychol. Rev. 65, 386–408 (1958).

40. Delépine, B., Duigou, T., Carbonell, P. & Faulon, J.-L. RetroPath2.0: A retrosynthesis workflow for metabolic engineers. Metab. Eng. 45, 158–170 (2018).

41. Delépine, B., Libis, V., Carbonell, P. & Faulon, J.-L. SensiPath: computer-aided design of sensing-enabling metabolic pathways. Nucleic Acids Res. 44, W226–31 (2016).

42. Perez, J. G., Stark, J. C. & Jewett, M. C. Cell-Free Synthetic Biology: Engineering Beyond the Cell. Cold Spring Harb. Perspect. Biol. 8, (2016).

43. Moore, S. J. et al. Rapid acquisition and model-based analysis of cell-free transcription-translation reactions from nonmodel bacteria. Proc. Natl. Acad. Sci. U. S. A. 115, E4340– E4349 (2018).

44. Karim, A. S., Heggestad, J. T., Crowe, S. A. & Jewett, M. C. Controlling cell-free metabolism through physiochemical perturbations. Metab. Eng. 45, 86–94 (2018).

45. de los Santos, E. L. C., Meyerowitz, J. T., Mayo, S. L. & Murray, R. M. Engineering Transcriptional Regulator Effector Specificity Using Computational Design and In Vitro Rapid Prototyping: Developing a Vanillin Sensor. ACS Synth. Biol. 5, 287–295 (2016).

46. Swank, Z., Laohakunakorn, N. & Maerkl, S. J. Cell-free gene regulatory network engineering with synthetic transcription factors. doi:10.1101/407999

47. Koch, M., Pandi, A., Delépine, B. & Faulon, J.-L. A dataset of small molecules triggering transcriptional and translational cellular responses. Data Brief 17, 1374–1378 (2018).

48. Libis, V., Delépine, B. & Faulon, J.-L. Expanding Biosensing Abilities through Computer-Aided Design of Metabolic Pathways. ACS Synth. Biol. 5, 1076–1085 (2016).

49. Cowles, C. E., Nichols, N. N. & Harwood, C. S. BenR, a XylS Homologue, Regulates Three Different Pathways of Aromatic Acid Degradation in Pseudomonas putida. Journal of Bacteriology 182, 6339–6346 (2000).

50. Nevozhay, D., Adams, R. M., Murphy, K. F., Josic, K. & Balázsi, G. Negative autoregulation linearizes the dose-response and suppresses the heterogeneity of gene expression. Proc. Natl. Acad. Sci. U. S. A. 106, 5123–5128 (2009).

51. Roquet, N. & Lu, T. K. Digital and analog gene circuits for biotechnology. Biotechnol. J. 9, 597–608 (2014).

52. Weiss, J. N. The Hill equation revisited: uses and misuses. The FASEB Journal 11, 835– 841 (1997).

53. Qian, Y., Huang, H.-H., Jiménez, J. I. & Del Vecchio, D. Resource Competition Shapes the Response of Genetic Circuits. ACS Synth. Biol. 6, 1263–1272 (2017).

54. Zucca, S., Pasotti, L., Mazzini, G., De Angelis, M. G. C. & Magni, P. Characterization of an inducible promoter in different DNA copy number conditions. BMC Bioinformatics 13 Suppl 4, S11 (2012).

55. Karig, D. K., Iyer, S., Simpson, M. L. & Doktycz, M. J. Expression optimization and synthetic gene networks in cell-free systems. Nucleic Acids Res. 40, 3763–3774 (2012).

56. Voyvodic, P.L., Pandi, A., Koch, M., Conejero, I., Valjent, E., Courtet, P., Renard, E., Faulon & J.L. and Bonnet, J. Plug-and-play metabolic transducers expand the chemical detection space of cell-free biosensors. Nat. Commun. 10, 1697 (2019).

57. Michel, E. & Wüthrich, K. Cell-free expression of disulfide-containing eukaryotic proteins for structural biology. FEBS J. 279, 3176–3184 (2012).

58. Oh, I.-S., Kim, D.-M., Kim, T.-W., Park, C.-G. & Choi, C.-Y. Providing an oxidizing environment for the cell-free expression of disulfide-containing proteins by exhausting the reducing activity of Escherichia coli S30 extract. Biotechnol. Prog. 22, 1225–1228 (2006).

59. Bishop, C. M. Pattern Recognition and Machine Learning. (Springer, 2016).

60. Haykin, S. O. Neural Networks and Learning Machines. (Pearson Higher Ed, 2011).

61. Jain, A. K., Jianchang Mao & Mohiuddin, K. M. Artificial neural networks: a tutorial. Computer 29, 31–44 (1996).

62. Weiss, R., Homsy, G. E. & Knight, T. F. Toward in vivo Digital Circuits. in Evolution as Computation (eds. Landweber, L. F. & Winfree, E.) 67, 275–295 (Springer Berlin Heidelberg, 2002).

63. Sarpeshkar, R. Analog Versus Digital: Extrapolating from Electronics to Neurobiology. Neural Comput. 10, 1601–1638 (1998).

64. Lewis, D. D., Villarreal, F. D., Wu, F. & Tan, C. Synthetic biology outside the cell: linking computational tools to cell-free systems. Front Bioeng Biotechnol 2, 66 (2014).

65. Noriega, Leonardo. Multilayer perceptron tutorial. *School of Computing*. Staffordshire University (2005).

66. Haykin, S. S. Neural Networks: A Comprehensive Foundation. (Upper Saddle River, N.J.: Prentice Hall, 1999).

67. Cybenko, G. Approximation by superpositions of a sigmoidal function. Math. Control Signals Systems 2, 303–314 (1989).

68. Rojas, R. Neural Networks: A Systematic Introduction. (Springer Science & Business Media, 2013).

69. Cherry, K. M. & Qian, L. Scaling up molecular pattern recognition with DNA-based winner-take-all neural networks. Nature 559, 370–376 (2018).

70. Duigou, T., du Lac, M., Carbonell, P. & Faulon, J.-L. RetroRules: a database of reaction rules for engineering biology. Nucleic Acids Res. 47, D1229–D1235 (2019).

71. Shin, J. & Noireaux, V. Efficient cell-free expression with the endogenous E. Coli RNA polymerase and sigma factor 70. J. Biol. Eng. 4, 8 (2010).

72. Sun, Z. Z. et al. Protocols for implementing an Escherichia coli based TX-TL cell-free expression system for synthetic biology. J. Vis. Exp. e50762 (2013).

73. Caschera, F. & Noireaux, V. Synthesis of 2.3 mg/ml of protein with an all Escherichia coli cell-free transcription-translation system. Biochimie 99, 162–168 (2014).

74. R: The R Project for Statistical Computing. Available at: https://www.r-project.org/.

75. RStudio. RStudio (2014). Available at: https://www.rstudio.com/.

## References

1. Daniel, R., Rubens, J. R., Sarpeshkar, R. & Lu, T. K. Synthetic analog computation in living cells. Nature 497, 619–623 (2013).

2. Libis, V., Delépine, B. & Faulon, J.-L. Expanding Biosensing Abilities through Computer-Aided Design of Metabolic Pathways. ACS Synth. Biol. 5, 1076–1085 (2016).

3. Cowles, C. E., Nichols, N. N. & Harwood, C. S. BenR, a XylS homologue, regulates three different pathways of aromatic acid degradation in Pseudomonas putida. J. Bacteriol. 182, 6339–6346 (2000).

4. parts.igem.org. Available at: http://parts.igem.org/Main_Page.

